# Viral evolution during primary infection in immunocompromised hosts

**DOI:** 10.1101/2025.09.01.673077

**Authors:** Morgan Craig, Xiaoyan Deng, David V. McLeod

## Abstract

The immune response to viral infection is a delicate balance. By perturbing this balance, immunodeficiencies are expected to influence within-host viral evolution. Indeed, the presence of immunocompromised hosts has been argued to be a source of novel viral variants in some infectious diseases, including SARS-CoV-2. However, these arguments rest upon between-host models and so the role of immunodeficiencies on within-host evolution in primary infections is poorly understood. Using a mechanistic immunological model, here we consider how different immunodeficiencies shape the orchestration of the immune response during primary infection. We study how this alters the viral fitness landscape, thus speeding and slowing viral evolution. We show that during acute infections, while immunodeficiencies in neutrophils and interferon initially speed viral evolution, by the time the infection is cleared, mutations are at lower frequencies than in immunocompetent hosts. In persistent infections, we show that while T cell deficiencies slow viral evolution, interleukin-6 and macrophage deficiencies speed viral evolution. Finally, we show that positive epistatic interactions arising due to the immunological response will accelerate the evolution of viral mutations affecting the ability of virions to evade different aspects of the immune response and to enter host cells.

## Introduction

The immune response to viral infection is tightly coordinated through a series of integrated, nonlinear networks^1^. Immunodeficiencies perturb this delicate balance, altering immunological responses and prolonging infection. In turn, immunodeficiencies alter the viral fitness landscape, influencing viral evolution and the speed of adaptation. Indeed, many viruses of importance for public health, including HIV^2^, influenza^3,4^, SARS-CoV-2^5,6^, and HSV^7^, show increased mutations and frequent within-host viral evolution in immunocompromised hosts^8,9^, depending on the type and severity of the immunodeficiency^10^.

Because immunocompromised hosts may shed virus throughout the course of infection, within-host evolution creates the possibility of forward transmission of novel viral mutations or variants. For example, several SARS-CoV-2 variants have been hypothesized to have originated from accelerated evolution in immunocompromised hosts^11-14^, in part due to their genetic differences from other variants circulating in the population^15^. Mathematical modelling of this hypothesis predicts that the longer duration of infection in immunocompromised hosts can speed viral evolution^16,17^ relative to transmission chains of acute infections, and can produce genetically differentiated variants in the presence of fitness valleys in the viral fitness landscape^18,19^.

However, by focusing on the between-host dynamics, this previous work ignores within-host processes. Mutations are treated as neutral within-host, with the fitness valley occurring because “intermediate” mutations are selected against during between-host transmission, before a “jackpot” mutation is acquired^18,19^. Recent within-host modelling, on the other hand, predicts the strength of immune pressure is critical to reproducing within-host evolutionary patterns^20^. Yet, as this work relies on statistical fitness landscapes that abstract the immunological dynamics, how the nature and severity of immunodeficiencies affect the speed of within-host viral adaptation remains poorly understood.

Here, we consider how immunodeficiencies affecting the orchestration of the immune response determine the speed of within-host viral evolution in acute and persistent infections. We show that during acute infections, immunodeficiencies in neutrophils and interferon have the largest evolutionary role: while they can speed evolution early in the infection, by the time the infection has been cleared, mutations will be at a lower frequency in immunocompromised hosts. During persistent infections, T cell concentrations play a pivotal role: T cell deficiencies slow viral evolution, while interleukin-6 and macrophage deficiencies overstimulate T cells, speeding viral evolution. We then show how immunological interactions between mutations affecting evasion of different aspects of the immune response can generate epistasis in fitness, further affecting the speed of viral evolution during persistent infections. Together, this work underlines how dysregulated immune responses impact within-host viral evolution, with potential consequences for individual and population health.

## Model

### Immune response to primary infection

We focus on an immunological model capturing the dynamics of a primary infection caused by a respiratory virus. In what follows, we highlight the key aspects of the model; more details are provided in the Supplementary Information 1, including information on parameter estimation. A model schematic is presented in Figure 1.

**Figure 1.**
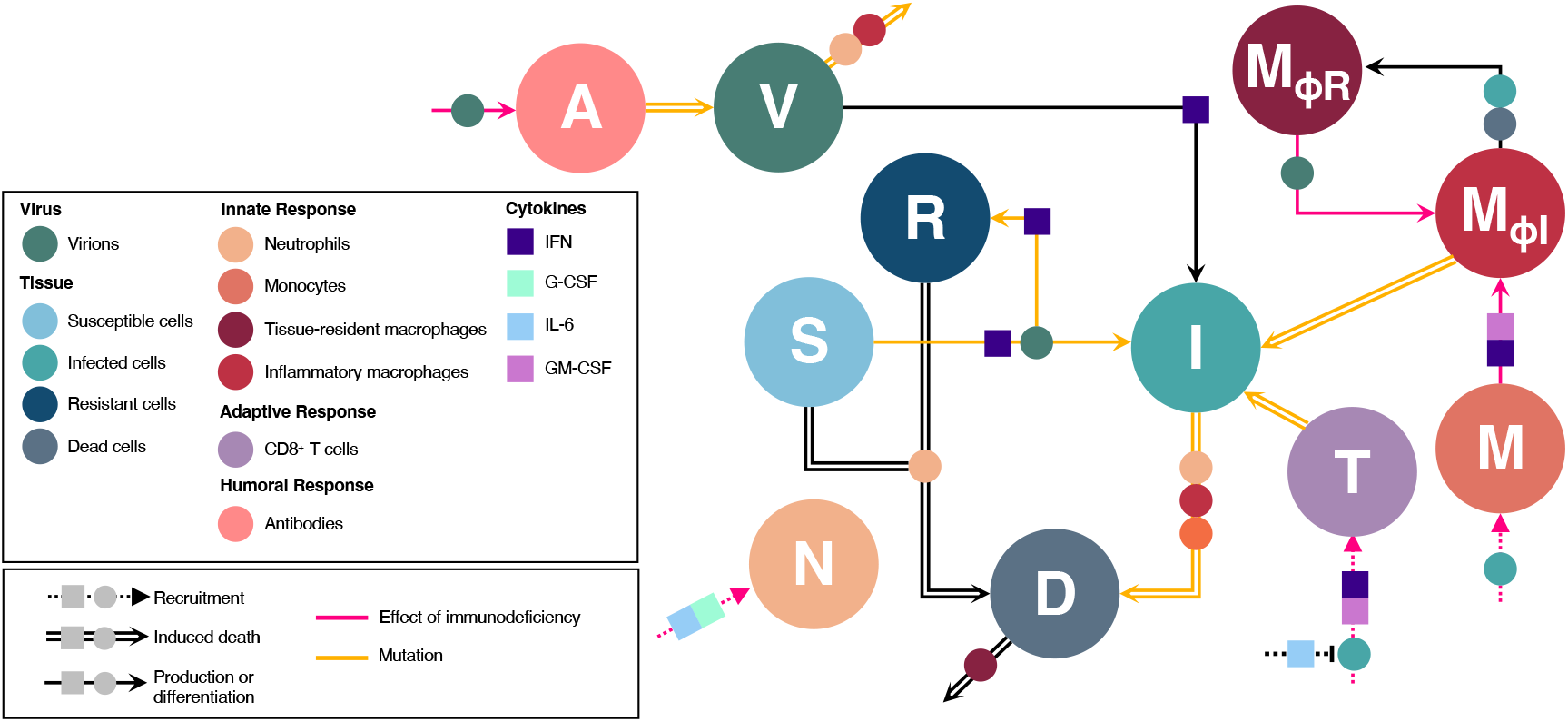
Schematic of immune response to viral infection. Virions, *V*, infect susceptible cells, *S*, producing infected cells, *I*. Following the eclipse phase, infected cells lyse, producing virions. Cells refractory to infection, *R*, are created through upregulation of interferon (IFN) signalling by infected cells. Inflammatory macrophages, *M*_Φ*I*_, are converted from monocytes, *M*, or tissue-resident macrophages, *M*_Φ*R*_, and along with neutrophils, *N*, phagocytose virions and infected cells. Phagocytosis of infected cells produces dead and damaged cells, *D*. Antibodies, *A*, are mobilized by the presence of viral antigens, and neutralize virions. CD8+ effector T cells, *T*, are stimulated by, and subsequently kill, infected cells. The action of the immune system is orchestrated through the antiviral cytokine IFN as well as the proinflammatory cytokines IL-6, G-CSF, and GM-CSF. Circles and boxes along arrows indicate the action of either a cell or cytokine, respectively. Dashed arrows: recruitment. Solid arrow: production or differentiation. Double arrows: induced death. Pink arrows: decreased effect due to immunodeficiency. Yellow arrows: immune effect targeted by mutations.

A primary infection consists of virions, *V*(*t*), and infected cells, *I*(*t*). Virions infect susceptible cells, *S*(*t*), via mass-action transmission with rate constant *β*. Infected cells undergo an eclipse phase lasting *τ*_*I*_ days during which cells are non-productively infected. Following the eclipse phase, lysis occurs with a burst size of *p* virions. Virions naturally decay at a per-capita rate *d*_*V*_, while the per-capita rate of infected cell death is *d*_*I*_.

Primary infection induces an immune response that can be divided into the innate, adaptive, and humoral responses. The innate immune response consists of three parts. First, following contact with virions, infected cells secrete Type I interferon (IFN) making them refractory to infection while also blocking viral entry and replication in neighbouring cells. Second, neutrophils, *N*(*t*), clear virions through neutrophil extracellular traps and kill infected cells. Neutrophils are recruited to the site of infection by signalling from the cytokines interleukin-6 (IL-6) and granulocyte colony-stimulator factor (G-CSF). Third, inflammatory macrophages, *M*_*ΦI*_(*t*), destroy virions and infected cells through phagocytosis^21^. Inflammatory macrophages are created from either monocytes, *M*(*t*), by signalling from the cytokines IL-6 and granulocyte macrophage colony-stimulator factor (GM-CSF), or from tissue-resident macrophages, *M*_*ΦR*_(*t*), after contact with cells that are infected, *I*(*t*), or dead, *D*(*t*).

The adaptive immune response consists of CD8+ effector T cells, *T*(*t*), and is activated *τ*_*T*_ days following viral infection. The activation delay accounts for the time necessary for naïve CD8+ T cells to be presented antigen, and to convert to the effector phenotype following a primary exposure^22^. Following activation, CD8+ effector T cells exhibit density-dependent saturable killing of infected cells^23^. CD8+ T cells are recruited by infected cells and IFN signalling and suppressed by IL-6, as IL-6 serves as a proxy for anti-inflammatory signalling^22^.

The humoral immune response consists of antibodies, *A*(*t*), which are generated by the presence of virions after *τ*_*A*_ days. The delay in antibody production accounts for the time required for antigen presentation to CD4+ T cells until the production of antibodies by B cells^24^. Following activation, antibodies reduce viral loads through saturable neutralization of virions. All antibodies are assumed to be neutralizing in our model.

Under these assumptions the dynamics of the virions and infected cells can be written

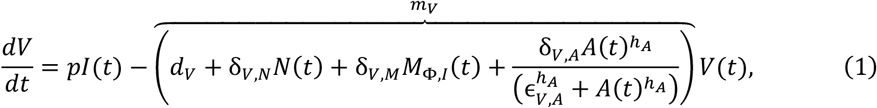

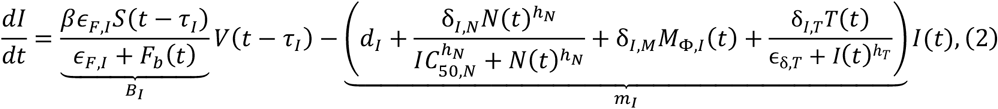

where *F*_*b*_(*t*) is the concentration of bound IFN at time *t*. In equations (1) and (2) we use *m*_*V*_ and *m*_*I*_ to denote the per-capita loss of virions and infected cells, respectively, and *B*_*I*_ to denote the per-capita production of infected cells by virions.

### Immunodeficiencies

Immunocompromised hosts may experience protracted “acute”, or persistent, infection and so experience a loss of IFN effects on refractory cells^25-28^, as well as a deficient innate, adaptive, and/or humoral immune response^29^. To capture the loss of IFN effects on refractory cells, we suppose they revert to susceptibility at a per capita rate *δ*_*R*_ after *τ*_*R*_ days. The delay in reversion accounts for the loss of increased Type I IFN signalling and interferon stimulated genes^30,31^ through e.g., receptor internalization and downregulation^32^.

To capture the deficient immune response, we suppose that the rate of production, and hence concentration, of different state variables of the immune response are reduced. For example, a monocyte production deficiency translates to a lower concentration of monocytes at homeostasis and during infection. The interconnectedness of the immune response means that production deficiencies in one variable may also impact other variables (e.g., the cytokine G-CSF recruits neutrophils, and so a G-CSF deficiency will also lead to a reduction in neutrophil concentrations). Therefore, for each immunodeficiency we first establish homeostasis at the specified level of production deficiency before introducing virions (see Sup. Info. Section 3 for more details).

We will assume immunodeficiencies reduce the target variable by at least 50% as compared to healthy hosts. For all state variables, except T cells and antibodies, this translates to reducing the production rate of the variable in question by 35% (Sup. Info. Section 3). To mimic clinically observed T cell immunodeficiencies, such as advanced HIV disease defined by CD4+ T cell counts <200 cells/*μl*^33^, a T cell production deficiency is assumed to reduce the production of T cells by 75%. To capture the near complete lack of antibodies in certain primary B cell immunodeficiencies^34^, including B cell lymphomas^8^, antibody immunodeficiencies are assumed to reduce antibody production by 90%. In all cases, varying the strength of the immunodeficiency quantitatively affects the reduction in the target variable, as well as the magnitude of concomitant changes in the non-target variables, but does not qualitatively affect our results (Sup. Info. Fig. S1).

### Viral evolution

As our interest is how immunodeficiencies shape (and speed) selection on viral evasion of the immune response, we ignore *de novo* mutations and assume that the viral mutation(s) of interest are present at a low frequency of 0.01 in the initial viral inoculum. Each viral mutation we consider targets evasion of a single immunological variable (e.g., neutrophils) in a particular life history stage (i.e., virion or infected cell). Each viral mutation is beneficial, and for simplicity, cost-free for the virus (the addition of costs is a straightforward extension). We will refer to the immunological variable directly targeted by the viral mutation as the “target” and the other immunological variables as “nontargets”.

The mutations considered can be divided into two groups, based on the life history stage (i.e., virion, infected cells) whose fitness they directly benefit. In the first group are mutations that directly increase a component of virion fitness (henceforth, *virion mutations*). Virion mutations target virion evasion of neutrophils (decreased δ_*V,N*_), macrophages (decreased δ_*V,M*_), or antibodies (decreased δ_*V,A*_), or target virion infection of susceptible cells, by increasing virion host cell entry^35,36^ (increased *β*), or by decreasing the ability of interferon to block viral entry^37,38^ (increased ϵ_*F,I*_). In the second group are mutations that directly increase a component of infected cell fitness (henceforth, *infected cell mutations*). Infected cell mutations target infected cell evasion of neutrophils (decreased δ_*I,N*_), macrophage (decreased δ_*I,M*_), or T cells (decreased δ_*I,T*_).

## Results

### The speed of within-host evolution of individual mutations

First, we ask how the immunological environment determines the speed of evolution of a single segregating mutation. To do so, we note that if *p*_*x*_(*t*) denotes the frequency of mutation *x* within the viral population, the frequency of mutation *x* changes according to

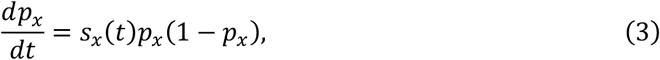

where *s*_*x*_(*t*) is the time-varying selection coefficient. Thus, the strength of selection on mutation *x* at time *t* is determined by the magnitude of *s*_*x*_(*t*); this dictates the instantaneous rate of increase of the mutation due to selection. In turn, *s*_*x*_(*t*) can be used to calculate the time-averaged strength of selection on mutation *x* from time *t*_0_ to time *t*:

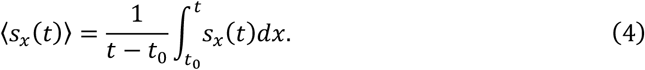

Equation (4) measures how strong constant selection would have to be to yield the observed change in mutation frequency by time *t*. Thus, *s*_*x*_(*t*) and ⟨*s*_*x*_(*t*)⟩ capture how the immune response determines the short- and long-term strength of selection, respectively, dictating the speed of adaptation, and therefore will be our focus here. Because the strength of selection will depend on the size of the mutational effect, in our figures we will normalize the selection coefficients, and the time-averaged strength of selection, by their maximum values over the duration of infection (e.g., *s*_*x*_(*t*)/max *s*_*x*_ (*t*)).

Calculating selection coefficients in populations with different life history stages (e.g., virions and infected cells) is not trivial, particularly for populations with temporally varying per-capita growth rates^39-42^. Therefore, we will primarily rely on two approximations for *s*_*x*_(*t*). First, we approximate *s*_*x*_(*t*) from simulation data as

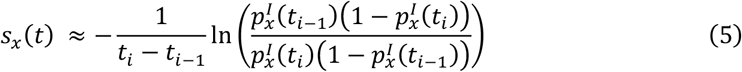

where 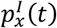 is the frequency of mutation *x* at time *t* in infected cells, and *t*_*i*_ − *t*_*i*−1_ is some (short) increment of time (e.g., hours, days). ⟨*s*_*x*_(*t*)⟩ can be approximated using equation (5) by setting *t*_*i*−1_ = *t*_0_. Although equation (5) only considers the frequency of the mutation in infected cells, simulation results indicate that it yields similar predictions if it is calculated using the frequency of the mutation in virions (Sup. Info. Fig. S2). The principal area of divergence between the different approximations of selection occurs at the outset of the infection. This is because each infection starts with only virions present, and so the delay in production of infected cells due to the eclipse phase means equation (5) cannot be accurately calculated during the initial few hours of the infection (Sup. Info. Fig. S2). Therefore, in all simulation results we plot equation (5) from hour 12 onwards.

Second, if we suppose mutation *x* is of weak phenotypic effect, and that the immunological variables, and density of susceptible cells, change slowly relative to the change in infected cells and virions (e.g., see Day et al.^39^), we can analytically approximate *s*_*x*_(*t*) as

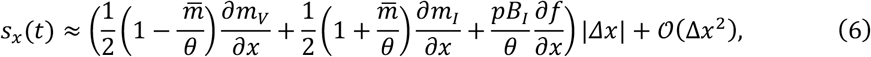

where |δ*x*| is the (small) phenotypic difference between mutant and wildtype, 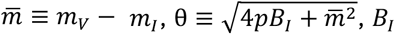 is evaluated when *τ*_*I*_ = 0, and *∂f*/*∂x* = 1 if *x* = *β* and *∂f*/*∂x* = *F*_*b*_/7ϵ_*F,I*_ + *F*_*b*_) if *x* = ϵ_*F,I*_ (Sup. Info. Section 4).

Equation (6) reveals that selection on each mutation is the product of two components. The first is the direct effect of mutation *x* on the viral fitness component *z* ∈ {*m*_*V*_, *m*_*I*_, *f*}, captured by 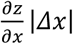. Since each mutation is beneficial and has a single effect on viral life history, only one of the direct effects in equation (6) will be nonzero and positive. The second component of selection in equation (6) is how each of the direct effects are weighted. These weights are based on the frequency of the mutation in the different life history stages (virions and infected cells) as well as the relative value of each life history stage for their contribution to future generations (reproductive value^39-42^).

As the per-capita rate of virion destruction is expected to be higher than that of infected cells, 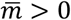, a decrease in *B*_*I*_ and/or *m*_*I*_ decreases the value of virion mutations and increases the value of infected cell mutations, whereas a decrease in *m*_*V*_ has the opposite effect. This has important consequences for the strength of selection.

### Availability of susceptible cells, and reversion to susceptibility of refractory cells, determines dynamical phases of primary infections

The availability of susceptible cells, and the reversion to susceptibility of refractory cells, partitions the infection into three phases exhibiting qualitatively distinct dynamics (Fig. 2). In the first phase of infection, susceptible cells are abundant and so the infection grows rapidly while the innate response (neutrophils, IFN, and macrophage) mobilizes (Fig. 2A-2B). As susceptible cells are abundant, the ratios 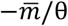 and *pB*_*I*_/θ are maximized. Thus, the value of virion mutations is at its maximum while the value of infected cell mutations is at its minimum (equation (6); Fig. 2G and 2H).

**Figure 2.**
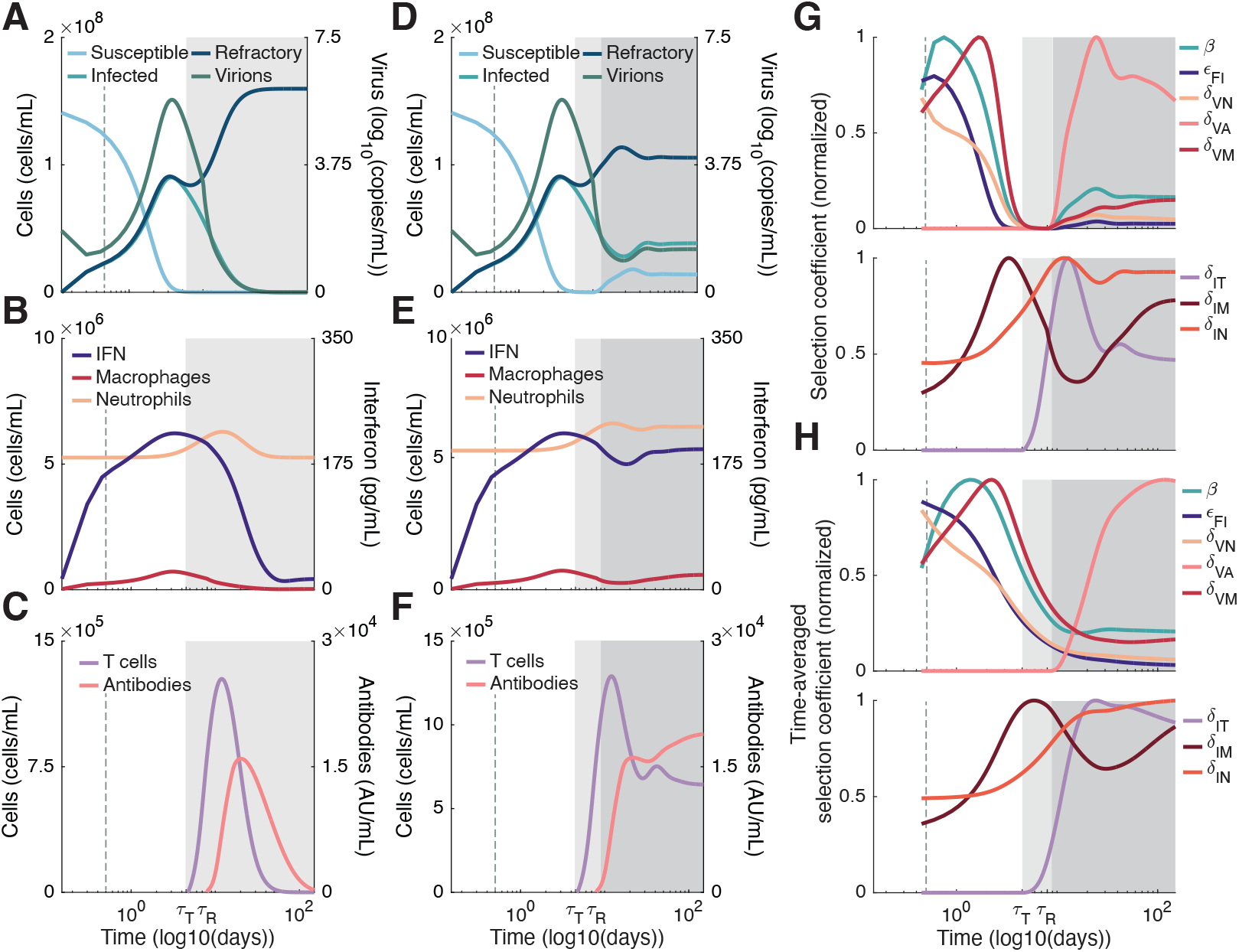
Immune dynamics distinguish the phases of infection. In immunocompetent hosts, the infection can be divided into two phases (panels A-C). In the first phase, indicated by the white area on each panel, susceptible cells are abundant, while the immune response is mobilizing. The abundance of susceptible cells means the value of virion mutations (i.e., mutations affecting cell entry *β*, as well as virion evasion of IFN, ϵ_*F,I*_, neutrophils, δ_*V,N*_, macrophage, δ_*V,M*_, and antibodies, δ_*V,A*_) is maximized, while the value of infected cell mutations (i.e., mutations affecting infected cell evasion of neutrophils, δ_*I,N*_, macrophage, δ_*I,M*_, and T cells, δ_*I,T*_) is minimized (panels G,H). At the same time, the mobilizing immune response (panels B,E) increases the direct effect of selection for immune evasion. In the second phase, indicated by light grey shading, susceptible cells are depleted, and the immune response eliminates the infection. During the second phase, the absence of susceptible cells means virion mutations are selectively neutral, while the value of infected cell mutations increases (panels G,H). In immunocompromised hosts (panels D-F), the reversion to susceptibility of refractory cells (δ_*R*_ > 0) leads to a third infection phase, indicated by dark grey shading on panels D-H. This causes a rebound in the value of virion mutations, and a decline in the value of infected cell mutations (panels G,H), while the mobilizing adaptive and humoral response (panels C,F) selects for T cell and antibody evasion. Note that in panels D-F we do not consider any additional immunodeficiencies other than δ_*R*_ > 0. In all panels, the dashed vertical grey line indicates 12 hours (see Sup. Info. Fig. S2).

In the second phase of infection, susceptible cells are largely depleted, and the innate immune response is fully mobilized. Because susceptible cells are depleted, *B*_*I*_ → 0, and so equation (6) reduces to

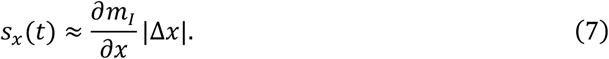

Thus, the value of virion mutations is zero and so virion mutations are selectively neutral, while the value of infected cell mutations is maximized (Fig. 2G and 2H).

In immunocompetent hosts (i.e., *δ*_*R*_ = 0), the depletion of susceptible cells combined with the mobilized immune response leads to clearance of the infection (Fig. 2A-2C). However, in immunocompromised hosts (Fig. 2D-2F), the loss of IFN signalling and reversion to susceptibility of refractory cells creates a third phase of infection shortly after *t* = *τ*_*R*_. In the third phase of infection, there is a temporary abundance of susceptible cells as the first wave of refractory cells revert to susceptibility. The dynamics then settle down to a quasi-equilibrium where the infection is limited by a combination of susceptible cell availability and the mobilized immune response (Fig. 2D-2F; dark grey shaded region). The rebound in susceptible cells in immunocompromised hosts increases the value of virion mutations and tends to decrease the value of infected cell mutations, as in the long term, *pB*_*I*_ → *m*_*V*_*m*_*I*_ (Fig. 2G,H). Moreover, the mobilization of the adaptive (T cells) and humoral (antibodies) response (Fig. 2C and 2F) generates a direct effect on selection for antibody and T cell evasion mutations, increasing the strength of selection on these mutations from the second to third phase of infection (Fig. 2G and 2H).

The time of onset of each phase of infection is affected by the size of the viral inoculum, *V*(0), in a predictable fashion. As the size of the inoculum decreases, the amount of time required for the viral population to exhaust susceptible cells increases, prolonging the first phase of infection, and so delaying the onset of the second phase (Sup. Info. Fig. S4). Although changes to the size of the viral inoculum affect the quantitative time of onset of the different phases, it does not qualitatively change the key predictions (Sup. Info. Fig. S4; see also Jenner et al.^22^). Therefore, in what follows we fix the initial inoculum at *V*(0) = 4.5 log_10_(copies/mL). For this inoculum size, the second and third phase of infection begin at approximately 4.5 and 9 days, respectively (Fig. 2D and 2F).

### Immunodeficiencies speed and slow the evolution of individual mutations

Next, we ask how different immunodeficiencies affect the speed of viral evolution. The most straightforward prediction occurs for target immunodeficiencies, that is, a production deficiency in the immunological variable directly targeted by the viral mutation. As the direct effect of mutation *x* on viral fitness component *z* ∈ {*m*_*V*_, *m*_*I*_, *f*} is an increasing function of the concentration of the target variable, target variable immunodeficiencies reduce the strength of selection on mutation *x* and so will slow the evolution of mutation *x*. This is intuitive: if, for example, mutation *x* targets antibody evasion and antibody concentrations are reduced, there are fewer antibodies to “evade” and so mutation *x* will be less beneficial, weakening selection and hence slowing evolution of mutation *x*. The interconnectedness of the immune response, however, means that non-target immunodeficiencies may not only feedback on the target variable, affecting the direct effect of unrelated mutations, but trigger changes in mutational value by, for example, affecting the availability of susceptible cells. In what follows, we consider these effects across the different phases of infection.

Because mutations targeting infected cell evasion of neutrophils (decreased *δ*_*I,N*_) and evasion of macrophage (decreased *δ*_*V,M*_ and *δ*_*I,M*_) are under very weak selection, regardless of the size of the mutational effect (Sup. Info. Fig. S2), in what follows we ignore these three mutations. We will therefore restrict our attention to mutations affecting cell entry (increased *β*), virion evasion of interferon, neutrophils, and antibodies (increased ϵ_*F,I*_ or decreased δ_*V,N*_ and δ_*V,A*_, respectively), and infected cell evasion of T cells (decreased δ_*I,T*_).

#### Neutrophil and IFN deficiencies tend to speed viral evolution in the first phase of infection, and slow viral evolution in the second phase

During the first two phases of infection, the immune response targeting infected cells and virions consists primarily of neutrophils, IFN, and macrophages (Fig. 2B,E). Of the three, neutrophils are the most important for controlling the initial infection (Fig. 3A). Neutrophils are recruited by the cytokine G-CSF; G-CSF is produced by monocytes; and monocytes are generated from bone marrow precursors by the cytokine GM-CSF (Fig. 1). A deficiency in any component of this pathway reduces neutrophil concentrations, either directly (i.e., a neutrophil deficiency) or indirectly (i.e., a G-CSF, GM-CSF, and/or monocyte deficiency), sharply increasing infection size (Fig. 3B and 3C). IFN deficiencies have a similar, but less strong effect. Since reduced neutrophil concentrations decrease the destruction of virions (decrease *m*_*V*_), while reduced IFN concentrations aid the infection of susceptible cells (increase *B*_*I*_), reductions in neutrophils and/or IFN increase the value of virion mutations and so tend to speed the evolution of non-target virion mutations targeting cell entry, neutrophil evasion, and IFN evasion during the first phase of infection (Fig. 3D and 3E, see also Sup. Info. Fig. S3A and S3H).

**Figure 3.**
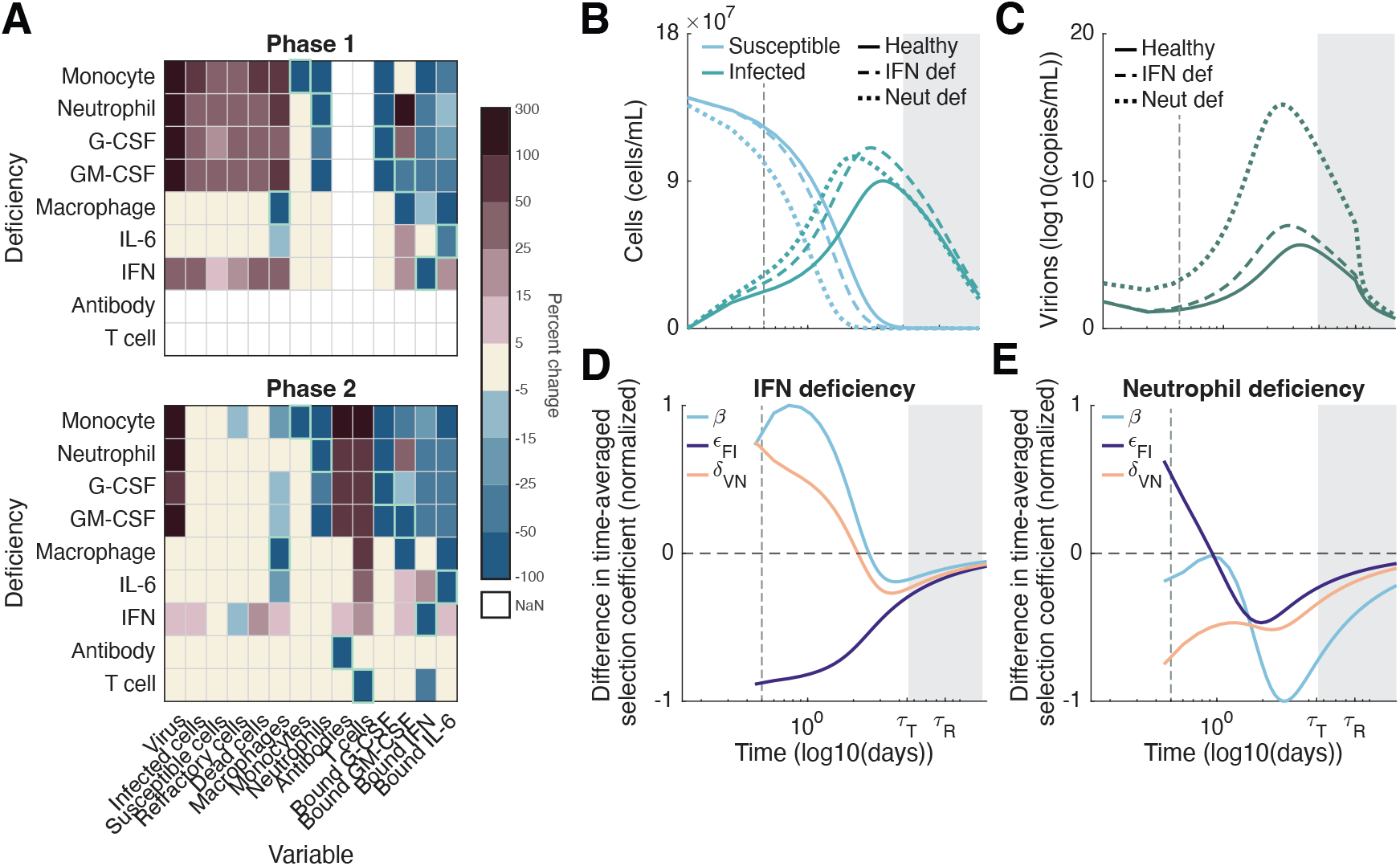
Viral evolution during acute infections is most strongly affected by neutrophil and IFN deficiencies. During acute infections (phase one and two; *δ*_*R*_ = 0), IFN and neutrophil deficiencies tend to speed evolution of non-target mutations during the first phase of infection, before slowing evolution during the second phase of infection (see also Sup. Fig. S3A and S3H). For acute infections, only mutations affecting cell entry, *β*, virion evasion of IFN, ϵ_*F,I*_, and virion evasion of neutrophils, δ_*V,N*_, experience significant selection. In panel A, the percent change in each immunological variable is calculated as (*X* − *Y*)/*X* × 100, where *X* and *Y* are the integrals of the indicated variable over the length of the phase in immunodeficient and healthy hosts, respectively, while the percent change in susceptible cells is calculated using the integral of *S*_*max*_ − *S*(*t*). In panels D and E, the difference in the time-averaged selection coefficient between immunodeficient and immunocompetent hosts is normalized by its maximum (absolute) value. The dashed vertical grey line in panels B-E indicates 12 hours.

However, reduced concentrations of neutrophils and/or IFN and the increase in infection size during the first phase of infection lead to a more rapid depletion of susceptible cells. This triggers an earlier onset of the second phase of infection. Since virion mutations are neutral during this phase (see equation (6)), by the end of the second phase of infection the normalized time-averaged selection coefficient is larger in immunocompetent hosts (Fig. 3D and 3E). Thus, if refractory cells do not revert to susceptibility (i.e., there is no third phase of infection), by the end of the infection, mutations are at a higher frequency in immunocompetent hosts.

#### T cell concentrations play a pivotal role in speeding or slowing viral evolution during phase three

If refractory cells do not revert to susceptibility, the infection is cleared during phase two. Therefore, to provide a point of comparison for the immunodeficiency of interest, in this section we allow for the loss of refractory cells (i.e., δ_*R*_ > 0) in both immunodeficient and immunocompetent hosts.

During the third phase of infection, T cells play an important role in killing infected cells, reducing the size of infection and increasing the availability of susceptible cells. This increases the value of virion mutations and decreases the value of infected cell mutations. T cell production is suppressed by bound IL-6 concentrations^43,44^; bound IL-6 is generated from stimulation of unbound IL-6 by neutrophils and monocytes, while unbound IL-6 is generated from infected cells, monocytes, and macrophages (Fig. 1; see also Sup. Info. 1). Consequently, while T cell deficiencies decrease T cell concentrations, deficiencies in neutrophils (arising either directly, or indirectly through deficiencies in G-CSF, GM-CSF, and/or monocytes), monocytes, macrophages and/or IL-6 will decrease bound IL-6 concentrations, increasing T cell concentrations (Fig. 4A and 4C), decreasing infection size and increasing the availability of susceptible cells (Fig. 4B). IFN and antibody deficiencies have limited effect on bound IL-6 or T cell concentrations (Fig. 4A).

**Figure 4.**
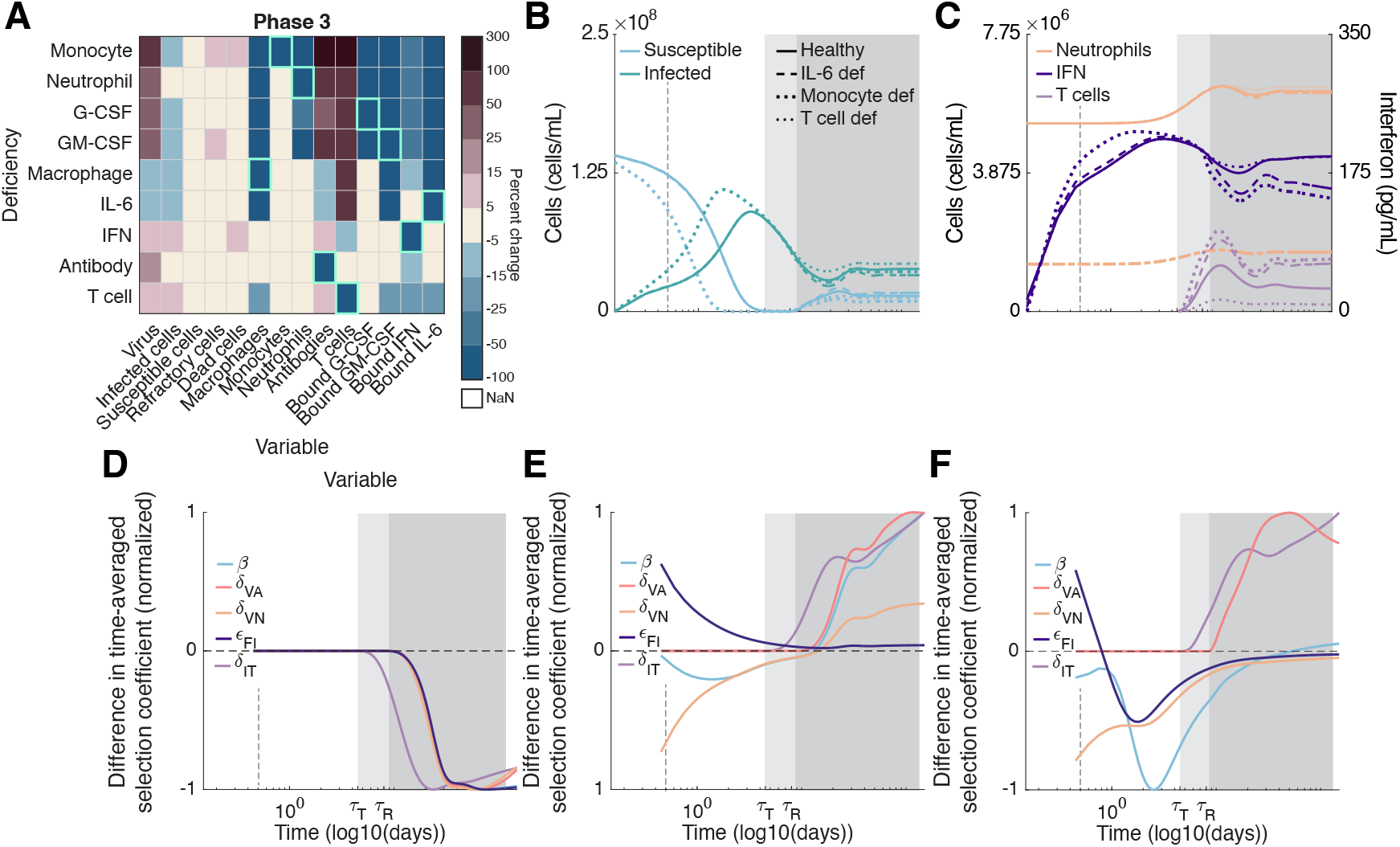
Immunodeficiencies affecting T cell concentrations strongly affect viral evolution during persistent infections. T cell killing of infected cells is a key determinant of the size of the infection and availability of susceptible cells (panels A and B) during persistent infections (i.e., *δ*_*R*_ > 0). A deficiency in T cells will increase infection size and decrease the availability of susceptible cells, decreasing the value of virion mutations and so slowing the evolution of all mutations considered (panel D). A deficiency in IL-6 (either directly, or due to a macrophage deficiency; Sup. Info. Fig. S3F) will increase T cell concentrations and so speed the evolution of all mutations considered (panel E). A deficiency in neutrophils (either directly, or due to a deficiency in the cytokines G-CSF and GM-CSF or monocytes, as shown here; see also Sup. Info. Fig. S3B-S3D), will increase T cell concentrations, but will also reduce neutrophils and IFN (panel F). In panel A, the percent change in each immunological variable is calculated as (*X* − *Y*)/*X* × 100, where *X* and *Y* are the integrals of the indicated variable over the length of the phase in immunodeficient and healthy hosts, respectively, while the percent change in susceptible cells is calculated using the integral of *S*_*max*_ − *S*(*t*). In panels D-F, the difference in the time-averaged selection coefficient between immunodeficient and immunocompetent hosts is normalized by its maximum (absolute) value. In panels B-F, the dashed vertical grey line indicates 12 hours.

Because a T cell deficiency both reduces the direct effect of T cell evasion as well as decreases the value of virion mutations by increasing susceptible cells, a T cell deficiency will slow the rate of increase of each of the mutations considered (Fig. 4F). On the other hand, since IL-6 and macrophage deficiencies increase T cell concentrations, they have the opposite effect, speeding the rate of increase of each of the five mutations considered (Fig. 4D and 4E). T cell counts are also increased by lower neutrophil concentrations arising through deficiencies in monocytes, G-CSF, GM-CSF, and/or neutrophils, as lower neutrophil concentrations will increase infection size. However, these deficiencies also reduce IFN (Fig. 4A). Consequently, deficiencies in monocytes, G-CSF, GM-CSF, and/or neutrophils speed the evolution of mutations targeting cell entry, antibody evasion, and T cell evasion, but slow the evolution of mutations targeting IFN and neutrophil evasion by reducing the concentration of IFN and neutrophils (Fig. 4D).

### Speed of evolution of multiple mutations

Finally, we ask how the speed at which individual mutations increase in frequency is affected by the presence of other segregating mutations. The possibility of genetic variation for multiple viral traits can yield fitness interactions between mutations (epistasis in fitness). Epistasis affects the speed of evolution by altering the fitness of a mutation depending on its genetic background, as well as producing linkage disequilibrium (LD) between mutations, that is, non-random assortment between mutations^45^; and the amount of LD in deterministic models will tend to scale with the strength of epistasis. For beneficial mutations, such as those studied here, these combined effects mean positive epistasis speeds evolution, whereas negative epistasis slows evolution^45-47^ (Sup. Info. 4.5).

Under the same assumptions that allowed us to derive equation (6), we can calculate a weak selection approximation of epistasis between pairs of mutations (Sup. Info. 4.5). From this approximation, two predictions emerge. First, if the two mutations both directly affect the fitness of virions (so virion evasion of neutrophils, antibodies, IFN, or increased cell entry) or both directly affect the fitness of infected cells, epistasis is positive, speeding their joint evolution (Sup. Info. 4.5). If instead, one mutation directly affects virion fitness, whereas the other mutation directly affects the fitness of infected cells (e.g., evasion of T cells), epistasis is negative, slowing their joint evolution (Sup. Info. 4.5).

Simulation results, however, suggest that while positive epistasis between virion mutations can dramatically speed evolution by enhancing fitness and by the generation of large amounts of LD, the ability of negative epistasis between virion and infected cell mutations to slow evolution is weak (Fig. 5). This occurs due to the source of epistasis, which depends upon the mutations under consideration. For example, mutations targeting cell entry and IFN evasion multiplicatively interact to affect the rate of infection of new cells, which will directly produce epistasis^48^. On the other hand, the source of negative epistasis in our model is strictly due to the division of the viral life cycle into virions and infected cells. This can be seen by supposing virion turnover is much more rapid than infected cell turnover, that is, *m*_*V*_ ≫ *m*_*I*_. As this difference grows, negative epistasis between virion and infected cell mutations becomes increasingly weak (Sup. Info. 4.5). Consequently, as our model parameterization assumes virion turnover is relatively rapid, negative epistasis is comparatively weak.

**Figure 5.**
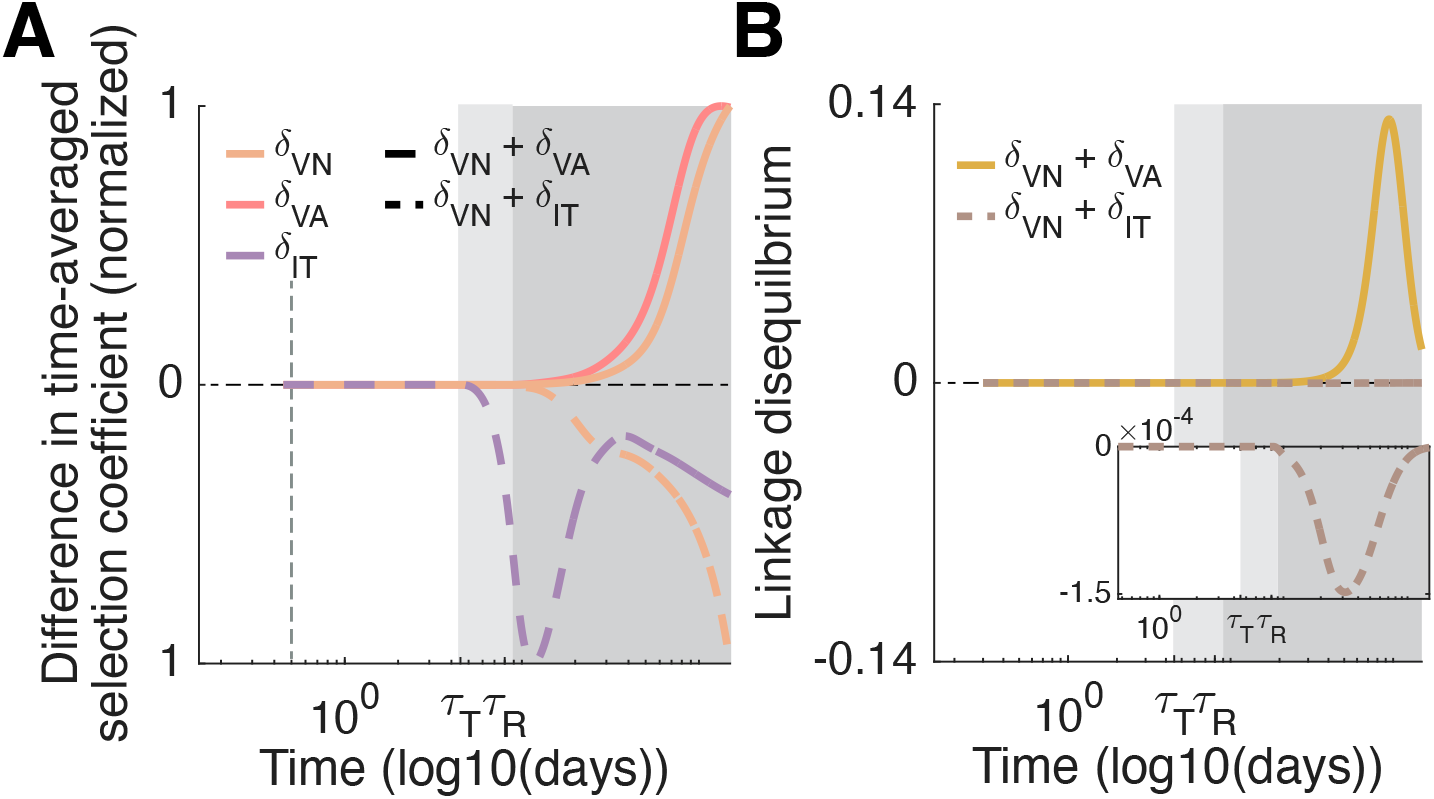
The sign of epistasis and the speed of viral evolution vary based upon the combination of mutations considered. When different mutations target immune evasion by the same life history stage, i.e., infected cells or virions, the speed of evolution can be dramatically increased through positive epistasis and the production of significant amounts of linkage disequilibrium (panels A and B, solid lines). Conversely, when one mutation targets T cell evasion, while the other targets virion evasion of the immune response, negative epistasis occurs (panels A and B, dashed lines). However, in contrast to positive epistasis, negative epistasis in our model is weak, and so is incapable of generating much linkage disequilibrium (panel B). That negative epistasis is weak arises due to the rapid turnover of virions relative to infected cells. In panel A, the difference in the time-averaged selection coefficients is normalized by their respective maximum (absolute) value.

## Discussion

Acute infections can become persistent in immunodeficient hosts due to weakened immune defences. Such persistent infections can provide an ideal environment for the generation of, and selection for, viral mutations^33,49^ increasing immune evasion^15,50,51^ and viral replication^10,52,53^. Indeed, between-host models have shown that persistent infections can speed viral evolution across the host population^16-19,54^. However, this previous work ignores how immunodeficiencies alter the within-host environment and the implications for viral evolution. Here, we work towards filling this gap by studying viral evolution during primary infection in immunocompromised hosts using a mechanistic model of the immune response.

The viral mutations considered in our model can be classified as “virion” or “infected cell” mutations, depending on the viral life stage they directly affect. The strength of selection for virion mutations tends to be positively correlated with the availability of susceptible cells and so follows a U-shaped dynamic through the phases of infection (Fig. 2G and 2H): virion mutations tend to be under strong selection at the outset of the infection when susceptible cells are abundant (phase one), they are selectively neutral once susceptible cells have been exhausted (phase two), and they rebound in value as refractory cells revert to susceptibility in persistent infections (phase three). Conversely, the strength of selection on infected cell mutations tends to be negatively correlated with the availability of susceptible cells and so follows an inverted U-shaped dynamic through the phases of infection.

The strength of selection on virion and infected cell mutations also depends on the type and severity of immunodeficiency. During an acute infection (phase one and two), immunodeficiencies in the orchestration of the innate immune response, specifically, deficiencies in IFN, neutrophils, monocytes and/or the cytokines G-CSF and GM-CSF play the most significant evolutionary role. Although such deficiencies strengthen selection on non-target virion mutations during the first phase of infection, selection is weakened during the second phase and by the end of an acute infection the frequency of virion mutations tends to be lower in hosts with these immunodeficiencies (Fig. 3D and 3E). During a persistent infection (phase three), T cell deficiencies increase infection size and so decrease the availability of susceptible cells. This slows the evolution of virion mutations, while also reducing selection on T cell evasion (Fig. 4D). Conversely, an IL-6 and/or macrophage deficiency overstimulates T cells, thus speeding the evolution of virion mutations and T cell evasion (Fig. 4E). While reduced neutrophil concentrations increase T cell counts, they also reduce IFN concentrations and so speed the evolution of mutations targeting cell entry, and antibody and T cell evasion, but slow the evolution of evasion of neutrophil and IFN (Fig. 4F).

Previous work using between-host models indicates that fitness valleys (i.e., negative epistasis), are necessary for immunocompromised hosts to promote the emergence of novel viral strains^18,19^. While fitness valleys are important at the between-host level, at the within-host level, it is probable positive epistasis is more likely to produce genetically distinct strains by accelerating evolution during persistent infections where viral diversity is more common. Indeed, our model indicates positive epistasis occurs between combinations of virion mutations and can significantly accelerate within-host evolution (Fig. 5). Conversely, negative epistasis, which occurs between virion mutations and mutations affecting T cell evasion, has a negligible effect (Fig. 5). Importantly, epistasis emerges in our model due to the immunological dynamics (i.e., from the phenotype-fitness mapping); previous work generated epistasis by building it into the genotype-phenotype map^18-20^.

Macrophages are an important component of the innate immune response, helping to bridge the innate and adaptive immune response^55^ and clear cellular debris^56^. As macrophage is primarily an orchestrator of the immune response, our model indicates evasion of macrophage is under weak selection (Sup. Info. Fig. S2) and macrophage deficiencies have limited evolutionary impact during acute infection. However, during persistent infections, macrophage deficiencies overstimulate T cells, speeding viral evolution. Mathematical modelling indicates macrophage deficiencies through dysregulated monocyte-to-macrophage differentiation, are predictive of disease severity^22^, as they cause T cell lymphopenia and hyperinflammation. Clinical studies have similarly reported shifts in the proportions of monocyte and macrophage subsets in the blood and lungs of severe COVID-19 patients, causing reductions to T cell recruitment of T cell and counts^57^. In combination with our analysis, this would suggest severe disease can also provide an optimal environment for within-host evolution.

To isolate how immunodeficiencies affect the orchestration of the immune response and the condition of the immunocompromised host at homeostasis, we intentionally constrained each immunodeficiency to reducing the production of a single immunological state variable. Real immunodeficiencies, however, involve multiple aspects of the immune response. For example, hosts with uncontrolled HIV show decreased production of naïve T cells, and CD4+ T cell lymphopenia^58^, resulting in a failure to produce antibodies^59^. Importantly, most clinically observed immunodeficiencies exhibit decreased T cells (see Sup. Info. 2.1), which our analysis reveals will slow viral evolution (Fig. 4D). A notable exception are hosts with B cell deficiencies and/or lymphomas^60^, who have weakened antibody responses and neutrophil deficiencies^34,61^; indeed, the latter speeds the evolution of T cell evasion (Fig. 4F).

In a secondary infection, the adaptive and humoral response will more rapidly activate leading to an earlier onset of selection for antibody and T cell evasion. Although beyond the scope of the current work, this raises the question of whether these mutations should be more strongly selected for in a primary infection in an immunocompromised host, or in a secondary infection in an immunocompetent host? Our analysis suggests that, owing to the heightened value of virion mutations at the beginning of the infection when susceptible cells are abundant, antibody evasion is more likely to be selected for in a secondary infection in an immunocompetent host. Conversely, as the value of infected cell mutations tends to be inversely correlated with the availability of susceptible cells, T cell mutations are more likely to experience stronger selection in a primary infection in an immunocompromised host.

To isolate the role of selection, we ignored the generation of *de novo* mutations, instead assuming mutations were present at low frequencies in the viral inoculum. It is well understood that, owing to the exponential within-host growth of viral populations during acute infections, most *de novo* mutations in acute infections will appear just before the susceptible cells are exhausted^62,63^, greatly restricting the amount of time for selection to operate. This constraint is alleviated in persistent infections, providing another reason for more rapid adaptation in immunocompromised hosts, as compared to chains of acute infections^63^. Furthermore, we assumed each mutation had a single beneficial effect on viral fitness and so ignored potential costs or trade-offs. Fitness trade-offs, however, are likely to affect the forward transmission process as within-host evolution is unlikely to optimize transmission between-hosts. Indeed, there is some data in SARS-CoV-2 evolution in immunocompromised hosts of a trade-off between antibody evasion and between-host transmissibility^53^. Thus, integrating our within-host evolutionary analysis into a between-host framework represents an important next step for future work to quantify the population-level risks posed by within-host evolution during persistent infections.

## Acknowledgements

The authors wish to thank the institutions who funded this study. MC is the Canada Research Chair in Computational Immunology and this research was undertaken, in part, thanks to funding from the Canada Research Chairs Program. MC was also funded by NSERC Discovery Grants RGPIN-2018-04546 and RGPIN-2025-04412. XD was funded by a Bourse de fin d’études from the Université de Montréal. DVM was funded by NSERC Discovery Grant RGPIN-2024-04608. MC and DVM were also funded by the NOVA – FRQNT-NSERC PROGRAM for junior researchers (https://doi.org/10.69777/342478). Funders played no role in study design, data collection, analysis and interpretation of data, or the writing of this manuscript.

## Supplementary Information

### 1. Immunological model of primary infection

Virions, *V*(*t*), infect susceptible cells, *S*(*t*), producing infected cells, *I*(*t*), via mass action transmission with rate constant *β*. Infected cells undergo an eclipse phase lasting *τ*_*I*_ days during which cells are non-productively infected. Following the eclipse phase, lysis occurs with a burst size of *p* virions.

Virions naturally decay at a per-capita rate *d*_*V*_ and are destroyed by neutrophils, *N*(*t*), inflammatory macrophages, *M*_*ΦI*_(*t*), and antibodies, *A*(*t*). Infected cells die at a per-capita rate *d*_*I*_, and are killed by neutrophils, inflammatory macrophages, and T cells, *T*(*t*). Infected cells can become refractory to infection, *R*(*t*), through Type-I interferon (IFN) signalling, *F*_*b*_(*t*). After *τ*_*R*_ time units, refractory cells revert to susceptibility at a per-capita rate *δ*_*R*_. The delay^30^ in reversion accounts for the loss of increased Type I IFN signalling and IFN stimulated genes^31^ through e.g., receptor internalization and downregulation^32^.

Susceptible and refractory cells are killed by neutrophil toxicity through bystander damage. These cells also undergo logistic growth, which depends on the concentration of susceptible, refractory, and infected cells, as well as the concentration of dead (and damaged) cells, *D*(*t*). Dead cells are generated by death or damage of infected, susceptible, and refractory cells. Dead cells naturally decay and are destroyed by macrophages.

Let

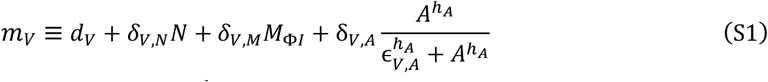

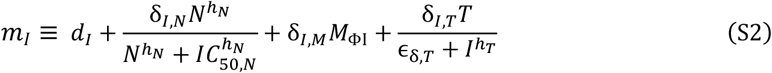

denote the per-capita decay rates of virions and infected cells, respectively. Then the dynamics of virions, and infected, susceptible, refractory, and dead cells are given by the following equations:

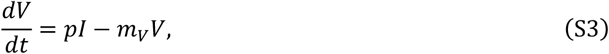

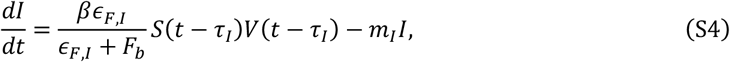

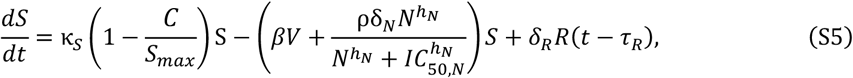

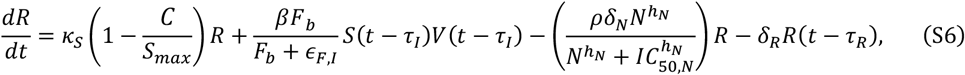

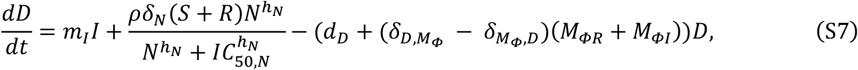

where *C* ≡ *S* + *I* + *R* + *D* is the total density of cells, regardless of state.

Infection first induces an innate response by neutrophils, *N*(*t*), and inflammatory macrophages, *M*_Φ*I*_(*t*). Neutrophils are generated by the bound cytokines interleukin-6 (IL-6; *L*_*b*_(*t*)) and granulocyte colony-stimulating factor (G-CSF; *C*_*b*_(*t*)) at a rate proportional to the concentration of neutrophils in the bone marrow, 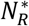, taken to be a constant for simplicity (here and in what follows we use a subscript *b* and *u* to denote bound and unbound cytokines). Neutrophils have a natural per-capita death rate of *d*_*N*_. Inflammatory macrophages are differentiated from monocytes, *M*(*t*), at a per-capita rate that depends on the cytokines IL-6 and granulocyte macrophage-colony stimulating factor (GM-CSF; *G*_*b*_(*t*)) given by

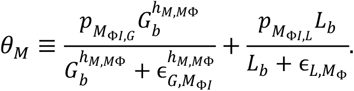

Inflammatory macrophages are also converted from tissue-resident macrophages, *M*_Φ*R*_(*t*), at a per-capita rate depending on infected and dead cells:

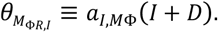

In turn, inflammatory macrophages convert to tissue-resident macrophages at a per-capita rate depending on tissue-resident macrophages and virions:

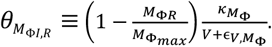

Macrophages are destroyed through phagocytosis and natural decay. Monocytes are generated by the cytokine GM-CSF at a rate proportional to the concentration of monocyte precursors in the bone marrow, 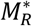 (taken to be a constant for simplicity). Monocytes are also generated through stimulation by infected cells. The dynamics of neutrophils, inflammatory and tissue-resident macrophages, and monocytes are given by

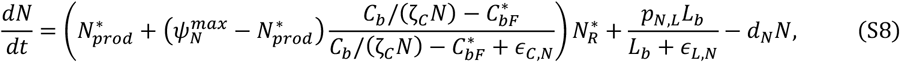

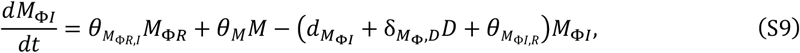

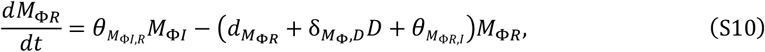

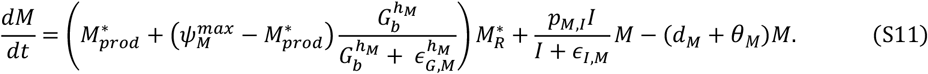

While the innate response is being mobilized, the humoral response mediated by antibodies, *A*(*t*), is generated by the presence of virions. The production of antibody by virions starts after *τ*_*A*_ days. The delay in antibody production accounts for the time required for the uptake of antigen by antigen presenting cells and their presentation to CD4+ T cells, the initial production of antibodies by short-lived plasma cells after CD4+ T cell priming, and the production of long-lived plasma cells and memory B cells from germinal centre B cells undergoing somatic hypermutation^24^. Antibodies neutralize virions and are lost due to natural decay. Their dynamics are given by

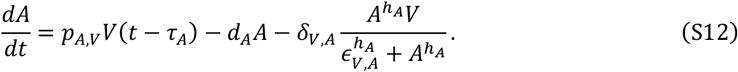

After *τ*_*T*_ days, the adaptive immune response mediated by CD8+ effector T cells, *T*(*t*), is activated. The activation delay accounts for the time necessary for naïve CD8+ T cells to be presented antigen, and to convert to the effector phenotype following a primary exposure^22,23,64,65^. CD8+ T cells are recruited by infected cells through a process mediated by IL-6^66^, are stimulated by IFN, and are lost due to natural decay. Their dynamics are given by

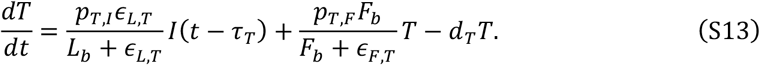

The action of the immune system is mediated by cytokines. We model unbound and bound cytokines, with bound cytokines responsible for immune effects. Let *Y*_*u*_(*t*) and *Y*_*b*_(*t*) denote the concentration of unbound and bound cytokine *Y* at time *t*, and *Y*_*prod*_(*t*) denote the rate of endogenous cytokine production. Then the general pharmacokinetic model of cytokine unbinding/binding is given by

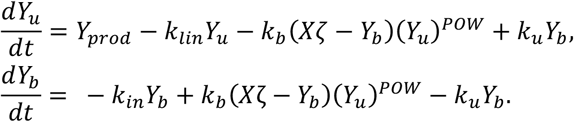

Here, *k*_*b*_ and *k*_*u*_ are the binding and unbinding rates, respectively, *k*_*in*_ is the rate of bound cytokine internalization, *k*_*lin*_ is the rate of elimination, *POW* is a stoichiometric constant, *X* is the sum of all cells modulated by the cytokine, and *ζ* is a scaling factor satisfying

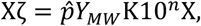

in concentration units of pg/mL. We used the equation *Y*_*MW*_ = *MM*/*N*_*A*_ to calculate the molecular weight of each cytokine, where *MM* is the molar mass and *N*_*A*_ = 6.02214 × 10^23^ is Avogadro’s number. In the equation above, 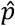 is a constant relating the stoichiometry between cytokine molecules and their receptors, *K* is the number of receptors specific to each cytokine on a cell’s surface and 10^*n*^ is a factor correcting for cellular units, giving:

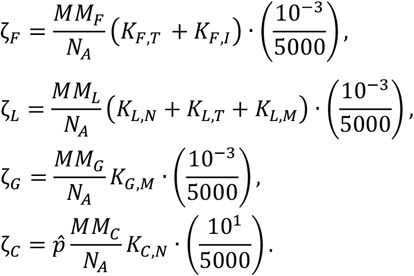

In our immunological model of primary infection, there are four key cytokines^22^: interferon (IFN), interleukin-6 (IL-6), granulocyte colony-stimulating factor (G-CSF), and granulocyte macrophage-colony stimulating factor (GM-CSF). Unbound IFN, *F*_*u*_(*t*), is produced by infected cells, monocytes, and inflammatory macrophages. IFN binds to infected cells and CD8+ T cells, producing bound IFN, *F*_*b*_(*t*), that leads to refractory cells. The dynamics of IFN are captured by

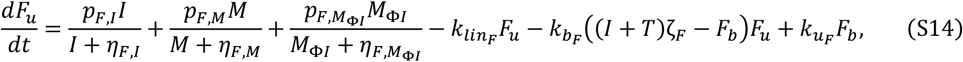

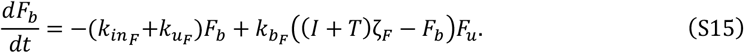

Unbound IL-6, *L*_*u*_(*t*), is produced by infected cells, monocytes, and inflammatory macrophages. IL-6 binds to neutrophils, monocytes, and CD8+ T cells, forming bound IL-6, *L*_*b*_(*t*), which stimulates neutrophil production and the differentiation of monocytes into inflammatory macrophage, and inhibits the production of CD8+ T cells. The dynamics of IL-6 are given by

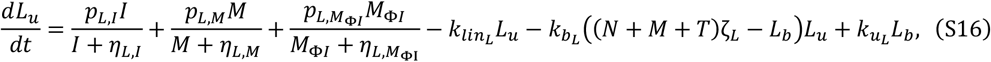

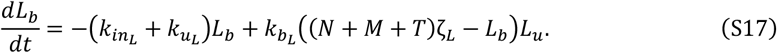

Unbound G-CSF, *C*_*u*_(*t*), is produced by monocytes. G-CSF binds to neutrophils, forming bound G-CSF, *C*_*b*_(*t*), which stimulates neutrophil production. The dynamics of G-CSF are given by

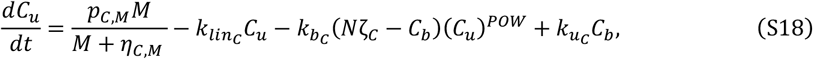

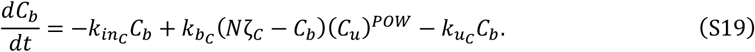

Finally, unbound GM-CSF, *G*_*u*_(*t*), is produced by monocytes and inflammatory macrophages and binds to monocytes, forming bound GM-CSF, *G*_*b*_(*t*). Bound GM-CSF produces monocytes and induces differentiation of monocytes into inflammatory macrophages. The dynamics of GM-CSF are given by

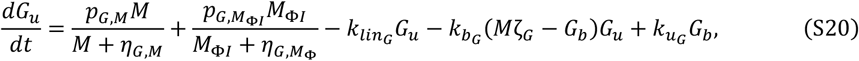

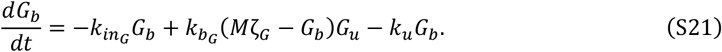

### 2. Immunological dynamics in immunocompromised hosts

Next, we modify the immunological model to allow for immunodeficiencies. We assume immunodeficiencies have two effects. First, they lead to prolonged infection due to the loss of IFN effects on refractory cells. To capture this, after *τ*_*R*_ time units, refractory cells revert to susceptibility at a per-capita rate *δ*_*R*_ in immunocompromised hosts (in immunocompetent hosts, *δ*_*R*_ = 0). Second, immunodeficiencies reduce the rate of production, and hence availability, of different aspects of the immune response. For example, a monocyte production deficiency translates to a lower concentration of monocytes at homeostasis and during infection.

Let *λ*_*i*_ denote a production deficiency in immune variable *i* ∈ {*N, M*_ϕ,*I*_, *A, T, F*_*u*_, *M, L*_*u*_, *C*_*u*_, *G*_*u*_}. Then production deficiencies modify the model as follows:

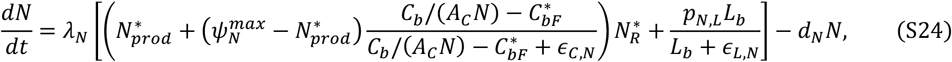

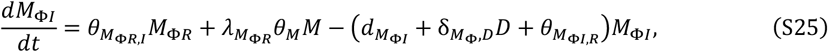

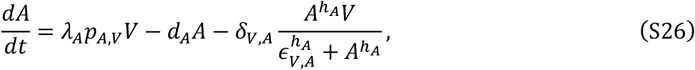

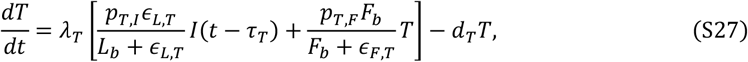

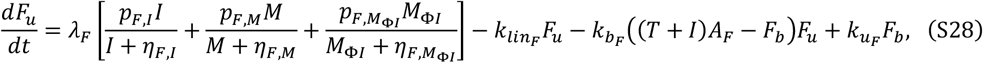

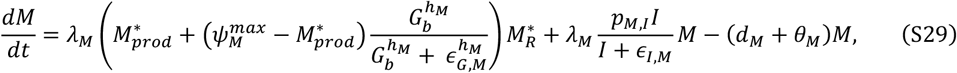

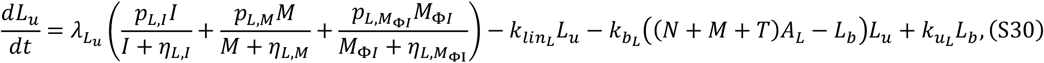

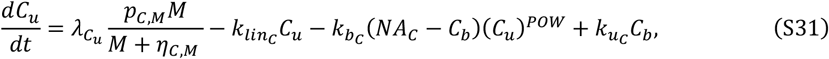

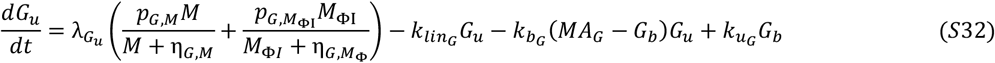

Production deficiencies also modify the initial conditions, as they can affect homeostasis in the absence of infection. Therefore, for all simulations we first allow the immune system to reach homeostasis before challenging the host with the viral inoculum.

#### 2.1 More realistic immunodeficiencies

While we treat immunodeficiencies as a reduction in the production rate of a single variable, clinically observed immunodeficiencies tend to be multifaceted, simultaneously affecting different aspects of the immune response. Four common immunodeficiencies, and their most prominent immunological features, are:

1. **B cell deficiencies and/or lymphomas**: These individuals have a weakened antibody response and may also experience neutrophil deficiencies^34,61^ (reduced *A*(*t*), *N*(*t*)), depending on the type of deficiency or lymphoma^60^.
2. **Uncontrolled HIV and AIDS**: Individuals with uncontrolled HIV show decreased production of naïve T cells, and CD4+ T cell lymphopenia^58^, resulting in a failure to produce antibodies^59^. Further, natural interferon-α producing cells are greatly reduced in individuals with AIDS^67^ (reduced *A*(*t*), *T*(*t*), *F*_*b*_(*t*))
3. **Non-B cell cancers**: Individuals with non-B cell malignancies are treated with myelosuppressive chemotherapies that suppress aspects of the innate immune system (e.g., deficiencies in neutrophils and macrophages^68^) as well as CD8+ T cells^69^ (reduced *N*(*t*), *T*(*t*), *M*_Φ,*I*_(*t*)).
4. **Solid organ transplant recipients**: These individuals are treated with lifelong immunosuppressive drugs^70^ and so are deficient in all aspects of the immune response (reduced *A*(*t*), *N*(*t*), *T*(*t*), *M*_Φ,*I*_(*t*)).

We expect that more realistic immunodeficiencies recapitulate the strongest effects we observed when considering single-variable immunodeficiencies, and so we do not consider them further here.

### 3. Parameter definitions and parameter values used in simulations

Our immunological model is based off a previous published model^22^; we have made several modifications to this model which we detail here. First, we allow for the reconversion of refractory to susceptible cells after *τ*_*R*_ days in immunocompromised hosts. The rate of this reconversion, δ_*R*_, was taken to be 0.05 days^-1^, and *τ*_*R*_ was set to 8 days to fall within the range of previous estimates^27^.

Second, consistent with studies from the lungs of influenza infected mice^23^, we modelled infected cell killing by T cells to be density dependent and saturable using the function

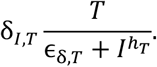

By comparing the total number of infected cells in our model versus that of Myers et al.^64^, we set the maximal rate of T cell killing to be δ_*I,T*_ = 7 days^-1^, the half-maximal cell concentration ϵ_*L,T*_ to be 0.01 x 10^9^ cells/mL, and the Hill coefficient *h*_*T*_ to be 0.5.

Third, given that primary infections can become protracted in immunocompromised hosts, we included neutralizing antibodies in our model. We assume antibodies are generated by the presence of virions *τ*_*A*_ days post-infection. The delay in antibody production accounts for the time required for the uptake of antigen by antigen presenting cells and their presentation to CD4+ T cells, the initial production of antibodies by short-lived plasma cells after CD4+ T cell priming, and the production of long-lived plasma cells and memory B cells from germinal centres^24^. We assumed antibodies are produced at a constant rate of *p*_*A,V*_ = 500 days^-1^ and are cleared at rate *d*_*A*_ = 0.033 days^-1^. We model the neutralization of virions by antibodies using the Hill function given by

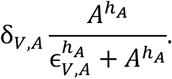

Based on our previous work where we fit this Hill function to data of SARS-CoV-2 neutralization by antibodies^24,71^, we set δ_*V,A*_ = 5 days-1, *h*_*A*_ = 1.19, and ϵ_*V,A*_ = 1000 AU/mL.

Fourth, we model immunodeficiencies as a reduction in the production of the target variable (see Sup. Info. 2). In our model, production of cells (i.e., neutrophils, monocytes, macrophages, and T cells) and cytokines (i.e., IFN, G-CSF, GM-CSF, and IL-6) is often generated through multiple pathways (see equations S24-S32). By assuming that immunodeficient hosts had at least a 50% reduction in the target variable compared to healthy hosts, we reduce the rate of production of neutrophils, monocytes, macrophages, IFN, G-CSF, GM-CSF, and IL-6 by 35%. We further reduce T cell and antibody production to more closely resemble clinical observations of lymphopenia^33^ and the absence of antibodies in certain patient groups^8,34^. Hence, T cell production is reduced by 75% and antibody production by 90%.

All other parameters in the immunological model were set to their previously estimated values. Full details are provided in the Supplementary Information to Jenner et al.^22^. Briefly, Jenner et al.^22^ applied a hierarchical estimation procedure in which certain parameter values were fixed directly from the literature, while others were fit using nonlinear least squares or nonlinear mixed effects methods to dose response data or time-series measurements.

In simulations incorporating immunodeficiencies, we calculated any remaining parameter values at physiological homeostasis to ensure that the immunological components return to basal concentrations. In particular, we set the differential equations for the different components of the immune system to 0 (in the absence of the virus), and then we calculated the values of 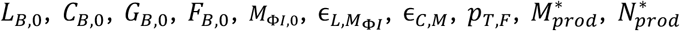, and 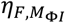. The complete list of parameter values is provided in Table S1.

**Figure S1.**
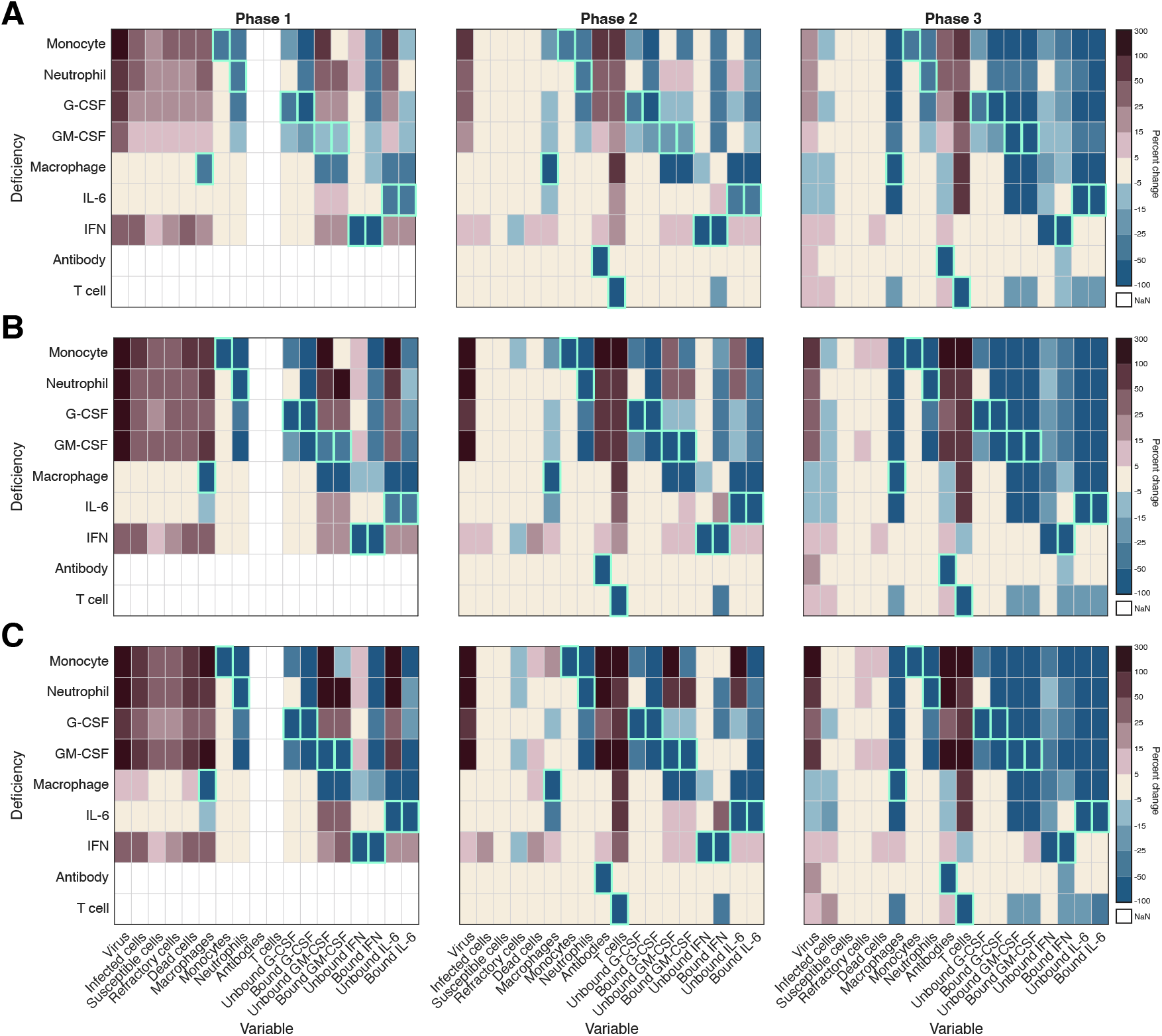
Changes in the immunological response in hosts with varying degrees of immunodeficiencies. Increasing the severity of the immunodeficiencies results in a quantitative, but not qualitative, change in the concentrations of the different immunological variables. From panel A to C, we increase the severity of the immunodeficiencies (panel B shows the severity of immunodeficiencies used in the main text). In panel A, the severity of immunodeficiencies was reduced by 25% from its baseline value shown in panel B (see Methods in the Main Text). In panel C, the severity of immunodeficiencies was increased by 25% from its baseline value (panel B). In all panels, the percent change in each immunological variable is calculated as (*X* − *Y*)/*X* × 100, where *X* and *Y* are the integrals of the indicated variable over the length of the phase in immunodeficient and healthy hosts, respectively, while the percent change in susceptible cells is calculated using the integral of *S*_*max*_ − *S*(*t*).

### 4. Viral evolution

#### 4.1 Evolutionary model

Next, we extend the immunological model to account for the possibility of viral evolution. To do so, we allow for strains carrying different mutations to simultaneously circulate in the population. We focus throughout on the action of selection and so ignore *de novo* mutations. We therefore assume that the mutant strains under consideration (and so the mutations of interest) are present at low frequencies (specifically, a frequency of 0.01) in the initial viral inoculum.

We consider eight possible mutations. Each of these mutations is beneficial and has a single effect on viral life history. The eight mutations considered are:

1. Virion evasion of antibodies, *A*(*t*), through decreased δ_*V,A*_.
2. Virion evasion of neutrophils, *N*(*t*), through decreased δ_*V,N*_.
3. Virion evasion of macrophages, *M*_Φ,*I*_(*t*), through decreased δ_*V,M*_.
4. Increased virion ability to enter the host cell, *S*(*t*), through increased *β* (e.g., by inhibiting PAMPs^35,36^).
5. Virion evasion of interferon, by reducing the ability of interferon to block viral entry through increased ϵ_*F,I*_.
6. Infected cell evasion of T cells, *T*(*t*), through decreased *δ*_*I,T*_.
7. Infected cell evasion of neutrophils, *N*(*t*), through decreased δ_*I,N*_.
8. Infected cell evasion of macrophages, *M*_Φ,*I*_(*t*), through decreased δ_*I,M*_.

The mutations can be grouped as either “virion mutations”, which are mutations that target components of virion fitness (mutations 1-5, that is, decreased δ_*V,N*_, δ_*V,M*_, δ_*V,A*_ or increased *β*, ϵ_*F,I*_), or “infected cell mutations”, which are the mutations that target components of infected cell fitness (mutations 6-8, that is, decreased δ_*I,N*_, δ_*I,M*_, δ_*I,T*_). Note that ϵ_*F,I*_ is classified here as a “virion mutation” mainly because the eclipse phase is short, and so it is more closely linked to virion fitness than infected cell fitness.

For each mutation, we divide the immunological variables into the variable directly targeted by the mutation (henceforth the “target”) and all other immunological variables not targeted by the focal mutation but targeted by a different mutation (henceforth, “nontargets”). For example, if the focal mutation targets virion evasion of neutrophils (decreased δ_*V,N*_), then the target variable is neutrophils, while the nontarget variables are macrophage, interferon, antibodies, T cells, and susceptible cells.

Next, let *I*_*i*_(*t*) and *V*_*i*_(*t*) denote the density of cells infected with strain *i* and strain *i* virions, respectively. Strain *i* has phenotype 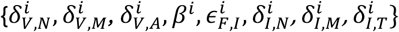. Then we can write the dynamics of strain *i* as

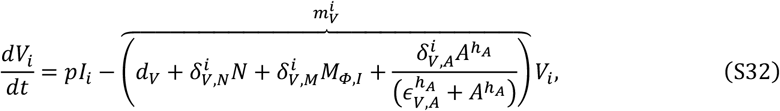

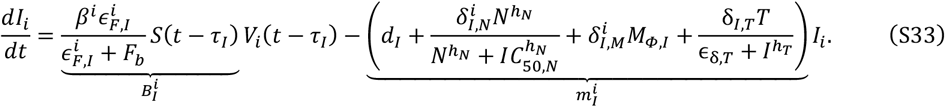

where 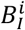 is the per-capita rate at which strain *i* virions produce cells infected by strain *i*, while 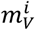 and 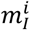 are the per-capita decay rates of strain *i* virions and infected cells, respectively.

#### 4.2 Evolution of individual mutations

First consider the evolution of each individual mutation. To do so, we note that if *p*_*x*_(*t*) denotes the frequency of mutation *x* within the viral population, the frequency of mutation *x* changes according to the equation

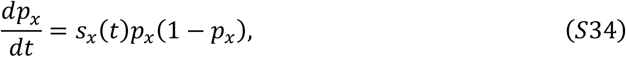

where *s*_*x*_(*t*) is the time-varying selection coefficient. The strength of selection on mutation *x* at time *t* is determined by the magnitude of *s*_*x*_(*t*); this dictates the instantaneous rate of increase of the mutation, given an observed mutation frequency. In turn, *s*_*x*_(*t*) can be used to calculate the time-averaged strength of selection on mutation *x* from time *t*_0_ to time *t*:

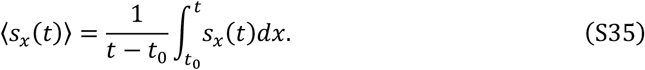

Equation (S35) measures how strong (constant) selection would have to be to yield the observed change in frequency by time *t*.

Thus, *s*_*x*_(*t*) and ⟨*s*_*x*_(*t*)⟩ capture how the immune response determines the short- and long-term strength of selection, respectively. This in turn dictates the speed of adaptation and so will be our focus here. Specifically, we will use *s*_*x*_(*t*) and ⟨*s*_*x*_(*t*)⟩ to understand how immunodeficiencies affect the speed of evolution of different mutations targeting viral evasion of the immune response.

##### 4.2.1 Approximating per-capita growth rate

To calculate the selection coefficient, we need to compute the per-capita growth rates, *r*_*i*_, of the different strains. This is because if there are two strains, *i* ∈ {*x*_*w*_, *x*_*m*_}, differing based on a single mutation, then the selection coefficient acting on mutation *x* is

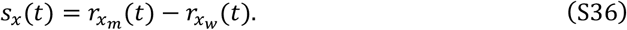

There are two challenges to computing per-capita growth rate in our model. The first challenge is the presence of the delay due to eclipse time, *τ*_*I*_. To remove the delay, define *E*_*i*_(*t*) to be the density of cells that have been infected by strain *i*, but are not yet productively infected. Then we can apply the approximation

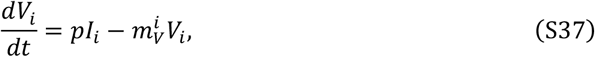

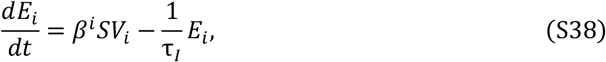

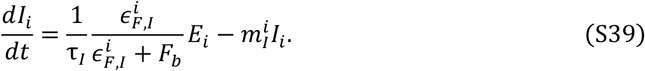

The second challenge to calculating per-capita growth rate arises due to the fact individuals belong to different classes (i.e., virions, infected cells, and productively infected cells). Calculating per-capita growth rates in populations where individuals belong to different classes (i.e., virions and infected cells) is not trivial, particularly for populations with temporally varying per-capita growth rates^39-42^. Therefore, to gain some analytic insight into how immunodeficiencies affect the strength of selection, suppose the immunological variables, and density of susceptible cells, change slowly relative to the change in infected cells and virions (e.g., see Day et al. 2022^39^). Then the per-capita growth rate of strain *i* is the dominant eigenvalue, *r*_*i*_, of the matrix

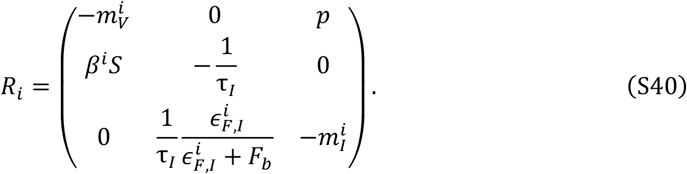

Because *R*_*i*_ is a 3 × 3 matrix, finding the dominant eigenvalue means finding the roots of the cubic polynomial:

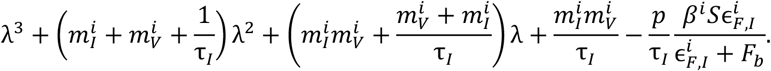

This is numerically straightforward but poses a challenge analytically. However, as the eclipse phase is short (τ_*I*_ is small), it is not unreasonable to suppose that the dynamics of non-productively infected cells occurs on a fast time scale, relative to the dynamics of virions and productively infected cells, that is *dE*_*i*_/*dt* ≈ 0, and so on the slow scale

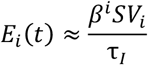

This approximation yields the reduced system on the slow time scale,

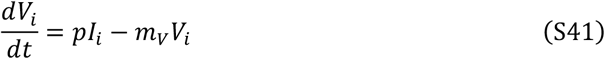

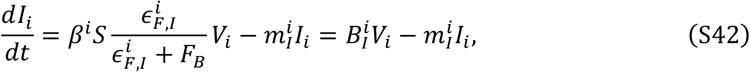

Where

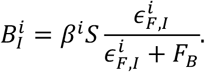

For the reduced system, the per-capita growth rate of strain *i* is the dominant eigenvalue, *r*_*i*_, of the matrix

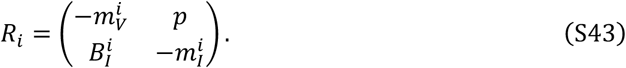

Thus

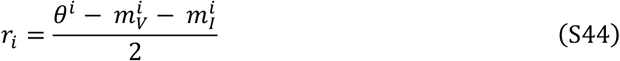

where 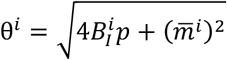 and 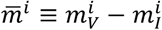. Numerical results indicate that approximating *r*_*i*_ using equation (S44) and using this to calculate the selection coefficient, *s*_*x*_, is very good, with the only deviation occurring at the beginning of the infection (Fig. S2).

For the numerical results presented in the main text, *s*_*x*_ can be approximated using simulation data as

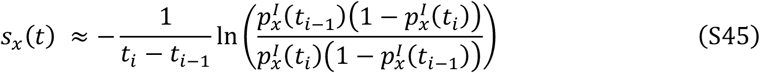

where 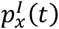 is the frequency of mutation *x* at time *t* in infected cells, and *t*_*i*_ − *t*_*i*−1_ is some (short) increment of time (e.g., hours, days). ⟨*s*_*x*_(*t*)⟩ can be approximated using equation (S45) by setting *t*_*i*−1_ = *t*_0_.

Although equation (S45) only considers the frequency of the mutation in infected cells, simulation results indicate that it largely yields similar predictions if it is calculated using the frequency of the mutation in virions (Fig. S2), even though the “true” selection coefficient involves a weighted average of the frequencies of virions and infected cells. The principal area of divergence between the different measures occurs at the onset of the infection. This is because each infection starts with only virions present, and so the delay in production of infected cells due to the eclipse phase means equation (S45) cannot be accurately calculated during the initial few hours of infection (Fig. S2). Moreover, the calculations of the selection coefficient using eigenvalues capture the strength of selection after the initial transient dynamics owing to the initial conditions are finished. Therefore, in all simulation results in the main text we plot equation (S45) from hour 12 onwards.

The other key point from Figure S2 is that mutations affecting virion and infected cell evasion of macrophage, and mutations targeting infected cell evasion of neutrophils are under very weak selection (Figure S2F-S2H). Specifically, the time for a mutation with a constant selection coefficient, *s*, to increase in frequency from *p*_0_ to *p*_1_ is

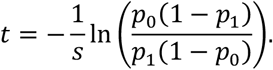

The time-averaged selection coefficient for mutations affecting evasion of macrophage and infected cell evasion of neutrophils are on the order of 10^−3^ or less, and so using the above equation it would take approximately 9,190 days for a mutation to increase from a frequency of *p*_0_ = 0.01 to *p*_1_ = 0.99. This very weak selection remains true regardless of the size of the mutational effect; indeed, in Figure S2F-S2H, the mutational effects for virion and infected cell evasion of macrophage, and infected cell evasion of neutrophils, are maximal.

The immunological reasoning for the negligible selection on mutations affecting δ_*V,M*_, δ_*I,M*_, and δ_*I,N*_ are as follows. The primary role of neutrophils during viral infections is to neutralize virions^72^ and not infected cells. Consequently, neutrophils have a saturated killing rate of infected cells^22^, which translates to weak selection on infected cell evasion of neutrophils. Macrophages, on the other hand, have two key effects. First, they balance inflammation and tissue repair, helping bridge the innate and adaptive immune responses^55^. Second, they clear pathogenic and cellular debris^56^ through phagocytosis once cells have become damaged or die. While the first effect is important for immune system functioning, it will not select for macrophage evasion, and the second effect occurs near the end of the infected cells life and so has limited selective consequences.

**Figure S2.**
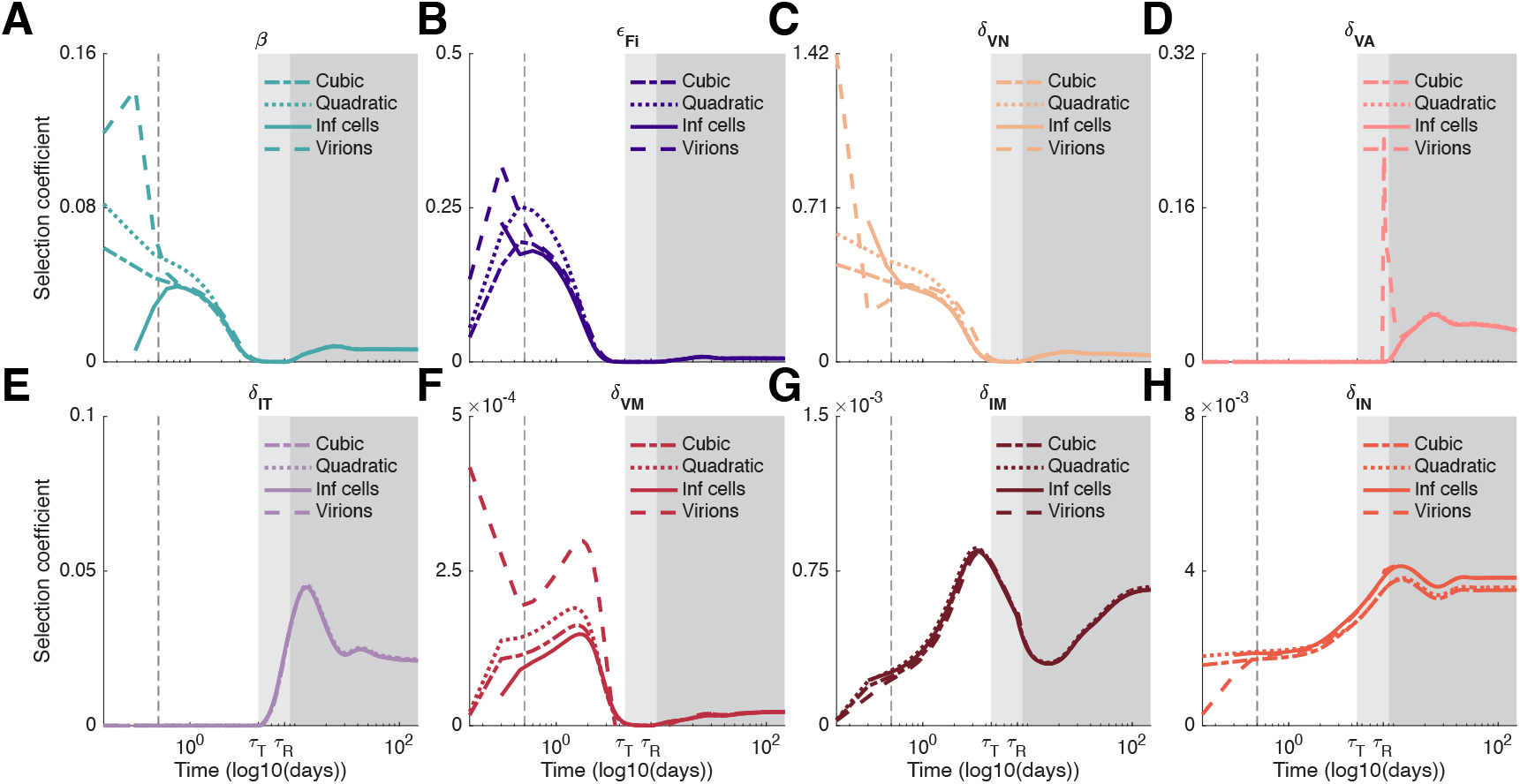
Comparison of di=erent approximations of the selection coe=icient. The selection coefficient can be numerically calculated in four ways: (1) by using formula (S36), with 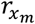 and 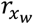 calculated using the matrix in equation (S40) (dashed-dot lines); (2) by using formula (S36), with 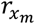 and 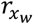 given by equation (S44) (dotted lines); (3) by applying equation (S45) to simulation data (solid lines); or (4) by applying equation (S45) to simulation data, except by measuring frequency amongst virions (dashed lines). Notice that virion and infected cell evasion of macrophage (panels F and G), and infected cell evasion of neutrophils (panel H) are under very weak selection, even though the mutational effect size is maximal. In all panels, the dashed vertical grey line indicates 12 hours.

##### 4.2.2 Weak selection approximation

We can gain further insight into the selection coefficient by supposing the mutational difference between the wildtype and mutant phenotypes, Δ*x* ≡ *x*_*m*_ − *x*_*w*_, is small, that is, selection is weak. Then we can use *r*_*i*_ to approximate *s*_*x*_ as

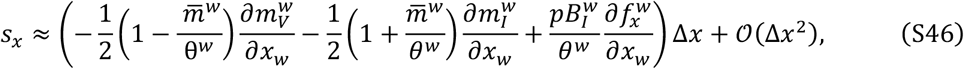

where *∂f*^*w*^/*∂x*_*w*_ = 1 if *x* = *β* and *∂f*^*w*^/*∂x*_*w*_ = *F*_*b*_/7ϵ_*F,I*_ + *F*_*b*_) if *x* = ϵ_*F,I*_.

Note that each of the *∂z*/*∂x*_*w*_ are non-negative for 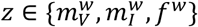, while δ*x* < 0 if mutation *x* affects the destruction of virions (*m*_*V*_) or infected cells (*m*_*I*_), whereas δ*x* > 0 if mutation *x* targets the infection of susceptible cells (*B*_*I*_). Consequently, each term in equation (S46) is positive, as observed in the main text.

From our approximation of *s*_*x*_, the strength of selection (magnitude of *s*_*x*_), depends on two factors. The first factor is the direct impact of mutation *x* on the relevant viral life-history quantity (i.e., the magnitude of 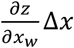, for *z* ∈ {*m*_*V*_, *m*_*I*_, *f*}. Since each mutation has a single effect on viral life history, only one direct effect will be non-zero. The second factor is how the direct effect is weighted due to class-structure (e.g., the magnitude of 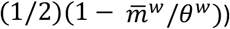. These weights capture the distribution of mutations between the different classes, as well as the reproductive value of each class.

In the first phase of the infection, *s*_*x*_ is as given in equation (S45). In the second phase of infection, susceptible cells have been largely depleted, that is, *S*(*t*) ≈ 0, and so 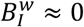.

Consequently, 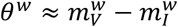 (since 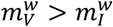), and so equation (S46) reduces to

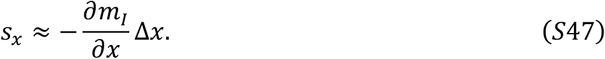

Thus, the value of virion mutations is zero, while the value of infected cell mutations is at its maximum. In the third phase of infection, eventually the concentration of virions and infected cells will be in a quasi-equilibrium state. Therefore, we have

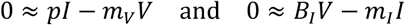

which implies

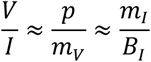

or *pB*_*I*_ ≈ *m*_*V*_*m*_*I*_. Consequently, θ ≈ *m*_*V*_ + *m*_*I*_, and so equation (S46) reduces to

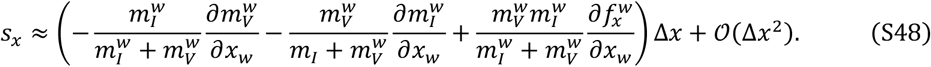

Because 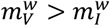, the value of mutations targeting infected cell evasion of the immune response is higher than the value of mutations targeting virion evasion of the immune response.

#### 4.3 Similar evolutionary dynamics are observed for di>erent immunodeficiencies

In the Main Text, we observed that certain immunodeficiencies have qualitatively similar effects on the strength of selection; this is shown in Figure S3. Immunodeficiencies in neutrophils, monocytes, and/or the cytokines G-CSF and GM-CSF induce qualitatively similar evolutionary dynamics (Fig. S3A-S3D). This is because each of these deficiencies trigger a reduction in neutrophil concentrations (Fig. S1). Similarly, immunodeficiencies in IL-6 and macrophage overstimulate T cell concentrations and so induce qualitatively similar evolutionary dynamics during persistent infections (Fig. S3E-S3F). T cell deficiencies and IFN deficiencies, on the other hand, yield distinct evolutionary dynamics (Fig. S3G-S3H).

**Figure S3.**
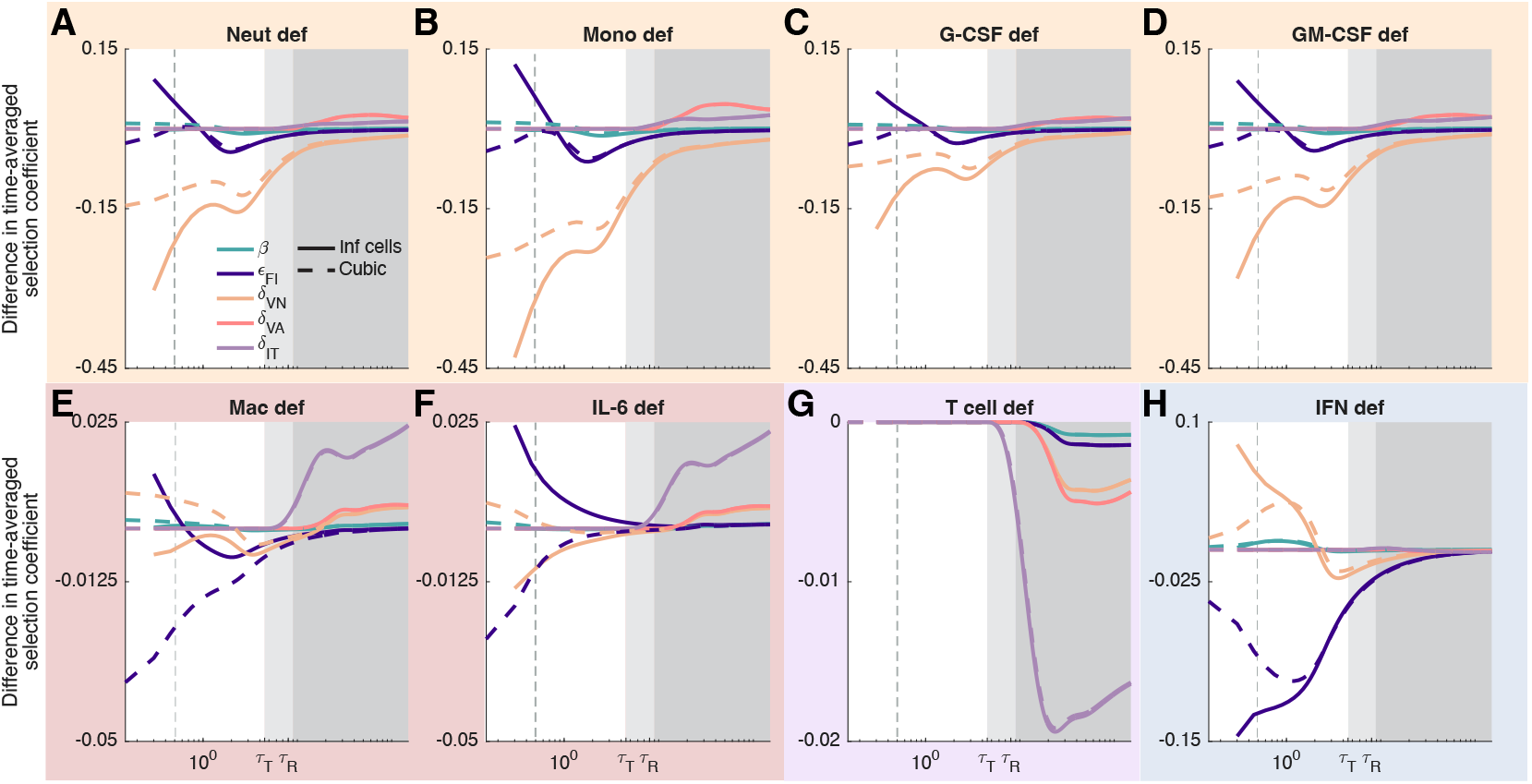
Qualitative similarities in evolutionary dynamics are determined by the type of immunodeficiency. Qualitatively similar viral evolutionary dynamics emerge for neutrophil, monocyte, G-CSF, and GM-CSF deficiencies (panels A-D, respectively; orange background), macrophage and IL-6 deficiencies (panels E and F, respectively; red background) while T cell (panel G; purple background) and IFN (panel H; blue background) deficiencies show distinct dynamics. In all panels, the time-averaged selection coefficient was calculated in two ways: using formula (S36), with 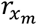 and 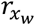 calculated as the dominant eigenvalue of matrix (S40) (dashed lines), or by applying equation (S45) to simulation data (solid line). The divergence between these measures at the beginning of the infection arises due to two sources: (1) estimating the selection coefficient from the simulation data only takes into account the frequency of mutations in infected cells, whereas the true selection coefficient is a weighted average of infected cells and virions, and (2) at the beginning of the infection, there are only virions present and there is a delay owing to the eclipse phase before infected cells appear. The dashed vertical grey line in each panel indicates 12 hours.

#### 4.4 Effect of inoculum size on infection dynamics

All infections initially consist of some concentration of virions, *V*(0) > 0, and no infected cells, *I*(0) = 0. However, we would expect some heterogeneity in the size of the initial inoculum across hosts. Heterogeneity in initial inoculum size affects the duration of phases of infection as well as the strength of the immune response. A smaller inoculum size slows the growth of the initial infection, slowing the depletion of susceptible cells and so extending the first phase of infection (Fig. S4). This will also tend to weaken the immune response across the first two phases of infection. The inoculum size has negligible consequences during the third phase of infection.

**Figure S4.**
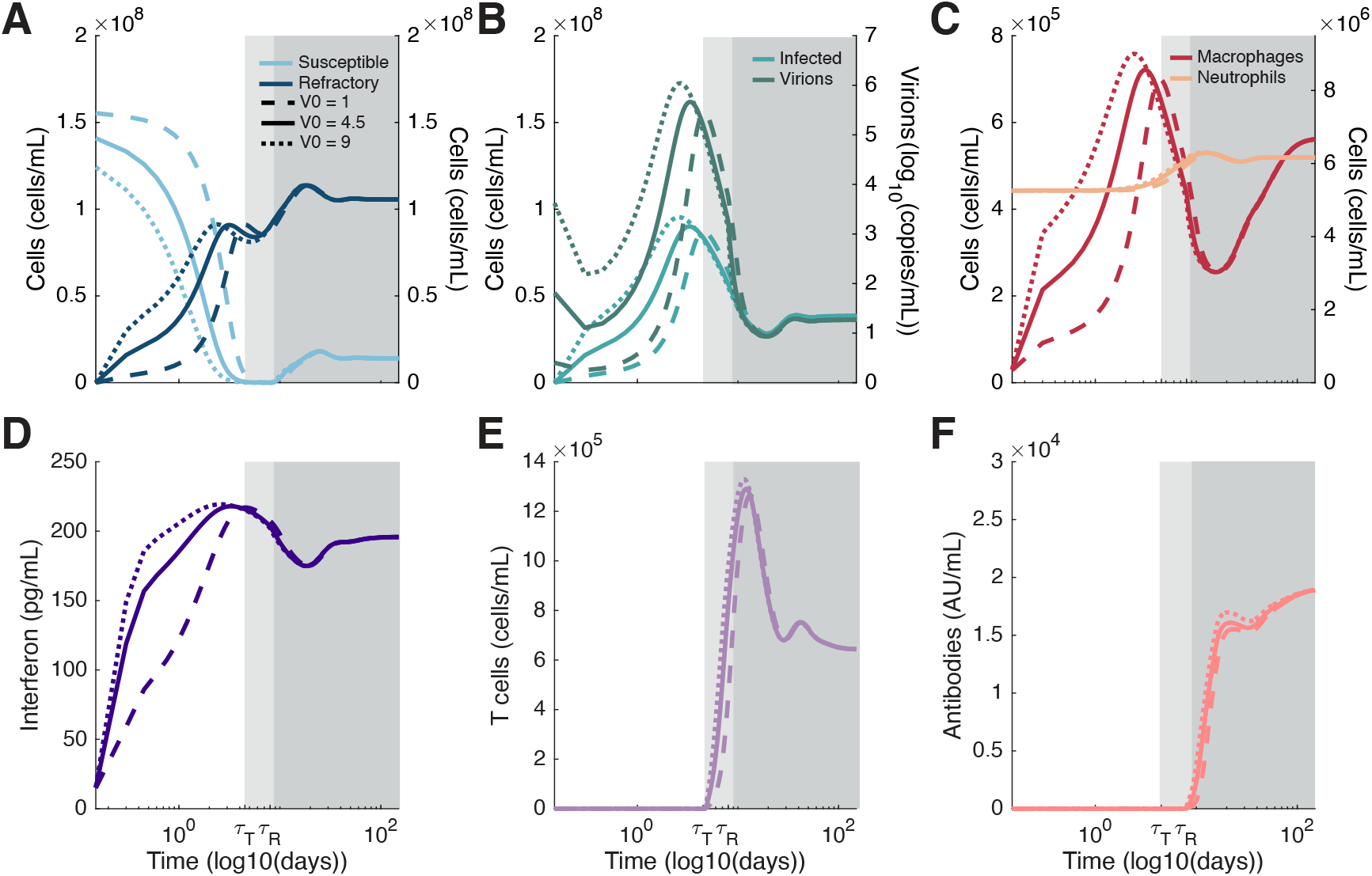
Inoculum size influences the onset of infection phases. Dynamics of susceptible and refractory cells (panel A), infected cells and virions (panel B), macrophages and neutrophils (panel C), interferon (panel D), T cells (panel E), and antibodies (panel F) for initial viral inoculum size of 1, 4.5, and 9 log_10_(copies/mL). Shaded regions indicate the three infection phases for an inoculum of 4.5 log_10_(copies/mL), as in the results shown in the main text. Larger inoculum sizes shorten (shift left) the first phase of infection due to higher viral loads and faster depletion of susceptible cells, while the reverse is true for smaller inoculum sizes.

#### 4.5 Multiple mutations

Next, we consider the evolution of multiple mutations. To do so, we suppose there are two mutations, *x* and *y*, at different loci affecting different aspects of the viral life history. This means there are four pathogen strains to consider, *i* ∈ {*x*_*w*_*y*_*w*_, *x*_*m*_*y*_*w*_, *x*_*w*_*y*_*m*_, *x*_*m*_*y*_*m*_}.

When mutations at different loci are simultaneously segregating in the population, in addition to selection on each individual mutation, we also need to account for epistasis in fitness between mutations. In continuous time models, if *r*_*i*_ is the per-capita growth rate of strain *i*, then epistasis in fitness is defined as

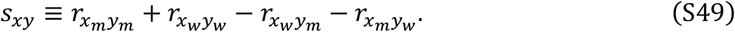

Thus, epistasis in fitness captures whether the fitness of a mutation depends on the genetic background or not.

If epistasis in fitness is positive, strains carrying both mutations are fitter than would be expected, based upon the fitness contribution of each individual mutation, speeding their evolution. If epistasis in fitness is negative, strains carrying both mutations are less fit than expected, slowing their evolution (in extreme cases, this can cause individually beneficial mutations to decrease in frequency). In addition, epistasis in fitness can produce linkage disequilibrium (LD) between mutations. The presence of LD, or the non-random associations between mutations, means that selection on mutation *y*_*m*_ will cause changes in the frequency of mutation *x*_*m*_ (so-called indirect selection). Because both mutations are individually beneficial, if epistasis is negative, then indirect selection will slow the evolution of both mutations, whereas if epistasis is positive, then indirect selection will speed the evolution of both mutations. Thus, in our model, indirect selection, and the direct consequences of epistasis work together to speed (positive epistasis) or slow (negative epistasis) evolution.

##### 4.5.1 Approximating epistasis in fitness

As before, per-capita growth rates are difficult to calculate in class-structured populations. Therefore, to make analytic progress we apply the same approximation as before. We assume that immunological variables, and density of susceptible cells, change slowly relative to the change in infected cells and virions and that the eclipse phase is short. Then under weak selection (i.e., the differences *x*_*m*_ − *x*_*w*_ and *y*_*m*_ − *y*_*w*_ are small), a Taylor series expansion of *s*_*xy*_ to leading order can be written

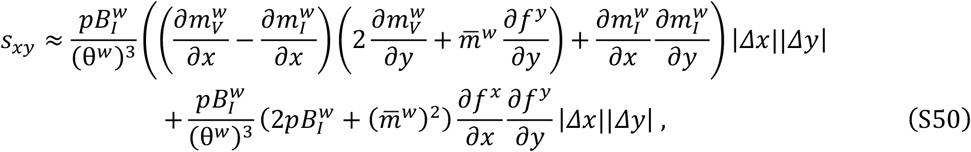

where *∂f*^*z*^/*∂z* = 1 if the mutation *z* affects transmissibility, *β*, and *∂f*^*z*^/*∂z* = *F*_*b*_/7*ϵ*_*F,I*_ + *F*_*b*_) if the mutation *z* affects evasion of IFN. As each mutation has a single direct effect, only one of the *∂q*/*∂x* and one of the *∂q*/*∂y* are non-zero for *q* ∈ {*m*_*V*_, *m*_*I*_, *f*}.

##### 4.5.2 The sign of epistasis in fitness

In our model, epistasis in fitness comes from two sources. First, if the two mutations target infection of susceptible cells (increased *β*) and interferon evasion (increased ϵ_*F,I*_), these two terms multiplicatively affect the production of infected cells. This produces a positive epistatic interaction affecting *B*_*I*_ that will ultimately translate to positive epistasis in fitness. Second, class structure means that even in the absence of epistatic interactions affecting one component of viral life history, epistasis in fitness tends to be generated^48^.

Recall that we previously identified two groups of mutations based on their direct fitness effects. In the first group are virion mutations. These mutations target virion infection and entry into cells (i.e., increased *β*, ϵ_*F,I*_) or target virion evasion of the immune response (i.e., decreased δ_*V,A*_, δ_*V,N*_, δ_*V,M*_). In the second group are infected cell mutations. These mutations target infected cell evasion of the immune response (i.e., decreased δ_*I,T*_, δ_*I,N*_, δ_*I,M*_). Our approximation of *s*_*xy*_ makes two predictions concerning these groups. If both mutations belong to the same group, epistasis in fitness is positive. If one mutation belongs to one group, and the other mutation belongs to the other group, epistasis in fitness is negative.

Negative epistasis is weak in our model. This occurs because the per-capita decay rate of virions is higher than the per-capita decay rate of infected cells. Consider the case in which 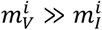. Then we can approximate the density of virions of strain *i* as

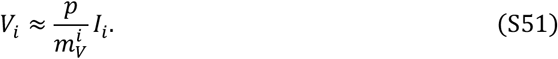

Using this relationship, we have

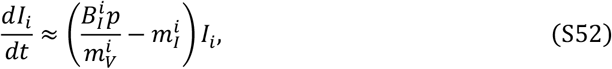

and so

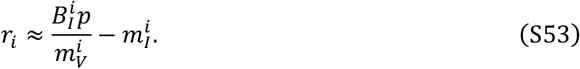

If we use this per-capita growth rate in the formula for epistasis given in equation (S49), we can see that negative epistasis is not possible between pairs of beneficial mutations.

**Table S1.** Parameter values for the immunological model.

| Parameter | Units | Description | Value |
| --- | --- | --- | --- |
| *Viral kinetic parameters* |  |  |  |
| $p$ | 1/day × log(cop/ml)<br>/10<sup>9</sup>cells | Lytic viral production | 2.59 |
| $\kappa_S$ | 1/day | Proliferation of epithelial cells | 0.7397 |
| $S_{max}$ | 10<sup>9</sup>cells | Epithelial cells carrying capacity | $S_0$ |
| $\kappa_{M\Phi}$ | log(cop/ml)/day | Production of alveolar macrophages | 5943 |
| $M_{\Phi max}$ | 10<sup>9</sup>cells/ml | Alveolar macrophage carrying capacity | $M_{\Phi R,0}$ |
| $\beta$ | 1/day × 1/<br>log(cop/ml) | SARS-CoV-2 virus infection rate | 0.30 |
| $\tau_I$ | days | Eclipse time | 0.17 |
| $\tau_T$ | days | Delay in CD8+ T cell arrival | 4.5 |
| $\tau_R$ | days | Delay in refractory cell re-entry | 8 |
| $\tau_A$ | days | Delay in antibody production | 5 |
| *Cell production, recruitment, and activation rates* |  |  |  |
| $p_{M\Phi I,G}$ | 1/day | Monocyte to macrophage differentiation<br>by GM-CSF | 1.68 |
| $p_{M\Phi I,L}$ | 1/day | Monocyte to macrophage differentiation<br>by IL-6 | 1.68 |
| $a_{I,M\Phi}$ | ml/(10<sup>9</sup>cells)×<br>(1/day) | Activation of macs by infected and dead<br>cells | $1.1 \times 10^3$ |
| $p_{M,I}$ | 1/day | Monocyte recruitment rate by infected<br>cells | 0.22 |
| $p_{T,F}$ | 1/day | CD8+ T cell production rate by IFN | 4 |
| $p_{N,L}$ | 1/day | Neutrophils recruitment rate by IL-6 | 0.3 |
| $p_{T,I}$ | 1/day | CD8+ T cell proliferation rate | 0.02 |
| $p_{A,V}$ | 1/day | Antibody production by virus | 500 |
| $M_{prod}^*$ | 1/day | Homeostasis reservoir release rate | 0.13 |
| $\psi_M^{max}$ | 1/day | Maximal reservoir release rate | 11.55 |
| $N_{prod}^*$ | 1/day | Homeostasis reservoir release rate | 0.21 |
| $\psi_N^{max}$ | 1/day | Maximal reservoir release rate | 4.13 |
| $C_{bF}^*$ | Dimensionless | Homeostasis neutrophil receptor bound<br>fraction | $1.6 \times 10^{-5}$ |
| *Cell-related half-effect ($\epsilon$), IC50 ($IC_{50}$), and Hill coefficient ($h$) parameters* |  |  |  |
| $\epsilon_{F,I}$ | pg/ml | IFN inhibition of viral production | $2 \times 10^{-4}$ |
| $\epsilon_{L,M\Phi}$ | pg/ml | IL-6 monocytes to macrophages | 1102.9 |
| $\epsilon_{G,M\Phi I}$ | pg/ml | GM-CSF monocyte to macrophages | 2664.5 |
| $\epsilon_{G,M}$ | pg/ml | GM-CSF recruitment of monocytes | 57.2 |
| $\epsilon_{F,T}$ | pg/ml | IFN production of CD8+ T cells | 399.3 |
| $\epsilon_{C,N}$ | unitless | G-CSF recruitment of neutrophils | $1.89 \times 10^{-4}$ |
| $\epsilon_{L,N}$ | pg/ml | IL-6 recruitment of neutrophils | 57.2 |
| $\epsilon_{I,M}$ | 10<sup>9</sup>cells/ml | Infected cell monocyte recruitment | 0.05 |
| $\epsilon_{L,T}$ | pg/ml | IL-6 production of CD8+ T cells | $1.5 \times 10^{-5}$ |
| $\epsilon_{V,M\Phi}$ | log(cop/ml) | Viral load for mac replenishing | 905.22 |
| $\epsilon_{\delta,T}$ | 10<sup>9</sup> cells/ml | Killing of infected cells by CD8+ T cells | 0.01 |
| $\epsilon_{V,A}$ | AU/ml | Antibody neutralization | 1000 |
| $h_M$ | Dimensionless | GM-CSF monocyte recruitment | 1.67 |
| $h_{M,M\Phi}$ | Dimensionless | GM-CSF monocyte to macrophages | 2.03 |
| $h_N$ | Dimensionless | Neutrophil induced damage | 3.02 |
| $h_A$ | Dimensionless | Antibody neutralization | 1.19 |
| $h_T$ | Dimensionless | Killing of infected cells by CD8+ T cells | 0.5 |
| $IC_{50,N}$ | 10<sup>9</sup>cells/ml | Neutrophil induced damage | $4.7 \times 10^{-5}$ |
| *Cell/virus-induced death rates* |  |  |  |
| $\delta_{V,M\Phi}$ | ml/(10<sup>9</sup>cells)×<br>1/day | Rate of viral clearance by macrophages | 768 |
| $\delta_{V,N}$ | ml/(10<sup>9</sup>cells)×<br>1/day | Rate of viral clearance by neutrophils | 1152 |
| $\delta_N$ | 1/day | Rate of neutrophil inflicted damage | 1.68 |
| $\rho$ | Dimensionless | Bystander death modulation constant | 0.5 |
| $\delta_{I,M\Phi}$ | ml/(10<sup>9</sup>cells)×<br>(1/day) | Rate macrophages phagocytose<br>infected cells | 121.20 |
| $\delta_{I,T}$ | ml/(10<sup>9</sup>cells)×<br>(1/day) | Rate CD8+ T cells induce apoptosis in<br>infected cells | 7 |
| $\delta_{M\Phi,D}$ | ml/(10<sup>9</sup>cells)×<br>(1/day) | Rate macrophages die from<br>phagocytosis | 6.06 |
| $\delta_{D,M\Phi}$ | ml/(10<sup>9</sup>cells)×<br>(1/day) | Rate macrophages phagocytose dead<br>cells | 8.03 |
| $\delta_R$ | 1/day | Rate of refractory cell reconversion | 0 (Immunocompetent)<br>0.05 (Immunodeficiency) |
| $\delta_{V,A}$ | log(copies/ml) x<br>(1/day) | Rate of antibody neutralization | 5 |
| *Cell death and decay rates* |  |  |  |
| $d_V$ | 1/day | Viral decay rate | 1.81 |
| $d_I$ | 1/day | Infected cell death rate | 0.1 |
| $d_D$ | 1/day | Degradation rate of apoptosed cells | 8 |
| $d_{M\Phi R}$ | 1/day | Alveolar macrophage death rate | 0.01 |
| $d_{M\Phi I}$ | 1/day | Inflammatory macrophage death rate | 0.3 |
| $d_M$ | 1/day | Monocyte death rate | 0.76 |
| $d_N$ | 1/day | Neutrophil death rate | 0.4 |
| $d_T$ | 1/day | CD8+ T cell death rate | 0.4 |
| $d_A$ | 1/day | Antibody clearance rate | 0.033 |
| *Cytokine production rates* |  |  |  |
| $p_{L,I}$ | pg/ml/day | IL-6 production by infected cells | 11.89 |
| $p_{L,M\Phi I}$ | pg/ml/day | IL-6 production by inflammatory<br>macrophages | 1872 |
| $p_{L,M}$ | pg/ml/day | IL-6 production by monocytes | 72.56 |
| $p_{G,M\Phi I}$ | pg/ml/day | GM-CSF production by inflammatory<br>macrophages | 2626 |
| $p_{C,M}$ | ng/ml/day | G-CSF production by monocytes | 26.26 |
| $p_{G,M}$ | pg/ml/day | GM-CSF production by monocytes cells | 123.4 |
| $p_{F,I}$ | pg/ml/day | IFN production by infected cells | 2.82 |
| $p_{F,M\Phi I}$ | pg/ml/day | IFN production by inflammatory<br>macrophages | 1.3 |
| $p_{F,M}$ | pg/ml/day | IFN production by monocytes | 3.56 |
| *Half-effect parameters* |  |  |  |
| $\eta_{L,I}$ | 10<sup>9</sup> cells/ml | IL-6 production by infected cells | 0.7 |
| $\eta_{L,M}$ | 10<sup>9</sup> cells/ml | IL-6 production by monocytes | 0.0045 |
| $\eta_{L,M\Phi I}$ | 10<sup>9</sup> cells/ml | IL-6 production by inflammatory<br>macrophages | 3.6× 10<sup>-5</sup> |
| $\eta_{G,M\Phi}$ | 10<sup>9</sup> cells/ml | GM-CSF production by macrophages | 3.6× 10<sup>-5</sup> |
| $\eta_{G,M}$ | 10<sup>9</sup> cells/ml | GM-CSF production by monocytes | 0.15 |
| $\eta_{C,M}$ | 10<sup>9</sup> cells/ml | G-CSF production by monocytes | 3.05 |
| $\eta_{F,I}$ | 10<sup>9</sup> cells/ml | IFN production by infected cells | 0.011 |
| $\eta_{F,M\Phi I}$ | $10^9$ cells/ml | IFN production by inflammatory macrophages | $1 \times 10^{-5}$ |
| $\eta_{F,M}$ | $10^9$ cells/ml | IFN production by monocytes | 0.54 |
| *Cytokine clearance and internalization rates* |  |  |  |
| $k_{linL}$ | 1/day | Rate of IL-6 renal clearance | 16.6 |
| $k_{linG}$ | 1/day | Rate of GM-CSF renal clearance | 11.7 |
| $k_{linC}$ | 1/day | Rate of G-CSF renal clearance | 0.16 |
| $k_{linF}$ | 1/day | Rate of IFN renal clearance | 18 |
| $k_{inL}$ | 1/day | Internalization rate of IL-6 | 61.8 |
| $k_{inG}$ | 1/day | Internalization rate of GM-CSF | 73.4 |
| $k_{inC}$ | 1/day | Internalization rate of G-CSF | 462 |
| $k_{inF}$ | 1/day | Internalization rate of IFN | 17 |
| *Cytokine binding/unbinding rates and stoichiometric constant* |  |  |  |
| $k_{bL}$ | ml/pg/day | IL-6 binding rate | 0.0018 |
| $k_{bG}$ | ml/pg/day | GM-CSF binding rate | 0.0021 |
| $k_{bC}$ | ml/ng/day | G-CSF binding rate | 2.24 |
| $k_{bF}$ | ml/pg/day | IFN binding rate | 0.011 |
| $k_{uL}$ | 1/day | IL-6 unbinding rate | 22.3 |
| $k_{uG}$ | 1/day | GM-CSF unbinding rate | 522 |
| $k_{uC}$ | 1/day | G-CSF unbinding rate | 184 |
| $k_{uF}$ | 1/day | IFN unbinding rate | 6.07 |
| $POW$ | Dimensionless | Stoichiometric constant (G-CSF) | 1.4608 |
|  |  | Stoichiometric constant (IL-6, GM-CSF, IFN) | 1 |
| $\hat{p}$ | Dimensionless | Stoichiometry relating constant (G-CSF) | 1.46 |
|  |  | Stoichiometry relating constant (IL-6, GM-CSF, IFN) | 1 |
| *Number of cellular receptors and cytokine molecular weights* |  |  |  |
| $K_{L,N}$ | sites/cell | No. IL-6 receptors on neutrophils | 720 |
| $K_{L,T}$ | sites/cell | No. IL-6 receptors on T cells | 300 |
| $K_{L,M}$ | sites/cell | No. of IL-6 receptors on monocytes | 509 |
| $K_{G,M}$ | sites/cell | No. GM-CSF receptors on monocyte | 1058 |
| $K_{C,N}$ | sites/cell | No. of G-CSF receptors on neutrophil | 600 |
| $K_{F,T}$ | sites/cell | No. of IFN receptors on T cells | 1000 |
| $K_{F,I}$ | sites/cell | No. of IFN receptors on infected cells | 1300 |
| $MM_L$ | g/mol | Molecular weight of IL-6 | 21000 |
| $MM_G$ | g/mol | Molecular weight of GM-CSF | 14000 |
| $MM_C$ | g/mol | Molecular weight of G-CSF | 19600 |
| $MM_F$ | g/mol | Molecular weight of IFN-β | 19000 |
| *Initial conditions* |  |  |  |
| $V_0$ | log(copies/ml) | Initial total viral load | 4.5 |
| $S_0$ | $10^9$ cells/ml | Initial susceptible cells | 0.16 |
| $I_0$ | $10^9$ cells/ml | Initial infected cells | 0 |
| $R_0$ | $10^9$ cells/ml | Initial refractory cells | 0 |
| $M_{\Phi R,0}$ | $10^9$ cells/ml | Initial resident macrophages | $2.73 \times 10^{-5}$ |
| $M_{\Phi I,0}$ | $10^9$ cells/ml | Initial inflammatory macrophages | $2.9 \times 10^{-7}$ |
| $M_0$ | $10^9$ cells/ml | Initial monocytes | 0.0004 |
| $M_R^*$ | $10^9$ cells/ml | Concentration of reservoir monocytes | 0.0023 |
| $N_0$ | $10^9$ cells/ml | Initial neutrophils | 0.0053 |
| $N_R^*$ | $10^9$ cells/ml | Concentration of reservoir neutrophils | 0.0316 |
| $T_0$ | $10^9$ cells/ml | Initial CD8+ T cells | $1.1 \times 10^{-4}$ |
| $L_{U,0}$ | pg/ml | Initial unbound IL-6 | 1.1 |
| $L_{B,0}$ | pg/ml | Initial bound IL-6 | 0 |
| $G_{U,0}$ | pg/ml | Initial unbound GM-CSF | 2.43 |
| $G_{B,0}$ | pg/ml | Initial bound GM-CSF | $1.6 \times 10^{-8}$ |
| $C_{U,0}$ | ng/ml | Initial unbound G-CSF | 0.025 |
| $C_{B,0}$ | ng/ml | Initial bound G-CSF | $6.5 \times 10^{-10}$ |
| $F_{U,0}$ | pg/ml | Initial unbound IFN | 0.015 |
| $F_{B,0}$ | pg/ml | Initial bound IFN | $1.1 \times 10^{-8}$ |
| $A_0$ | AU/ml | Initial antibodies | 0 |

## REFERENCES

1. Agrati, C. et al. Emerging viral infections in immunocompromised patients: A great challenge to better define the role of immune response. Frontiers in Immunology 14 (2023). 10.3389/fimmu.2023.1147871

2. Novak, R. M. et al. Prevalence of Antiretroviral Drug Resistance Mutations in Chronically HIV-Infected, Treatment-Naive Patients: Implications for Routine Resistance Screening before Initiation of Antiretroviral Therapy. Clinical Infectious Diseases 40, 468–474 (2005). 10.1086/427212

3. Renaud, C. et al. H275Y Mutant Pandemic (H1N1) 2009 Virus in Immunocompromised Patients. Emerging Infectious Diseases 17, 653–660 (2011). 10.3201/eid1704.101429

4. Weinstock, D. M., Gubareva, L. V. & Zuccotti, G. Prolonged Shedding of Multidrug-Resistant Influenza A Virus in an Immunocompromised Patient. New England Journal of Medicine 348, 867–868 (2003). 10.1056/nejm200302273480923

5. Raharinirina, N. A. et al. SARS-CoV-2 evolution on a dynamic immune landscape. Nature 639, 196–204 (2025). 10.1038/s41586-024-08477-8

6. Wagner, C. et al. Positive selection underlies repeated knockout of ORF8 in SARS-CoV-2 evolution. Nature Communications 15 (2024). 10.1038/s41467-024-47599-5

7. Gaudreau, A., Hill, E., Balfour, H. H., Erice, A. & Boivin, G. Phenotypic and Genotypic Characterization of Acyclovir-Resistant Herpes Simplex Viruses from Immunocompromised Patients. Journal of Infectious Diseases 178, 297–303 (1998). 10.1086/515626

8. Fournelle, D. et al. Intra-Host Evolution Analyses in an Immunosuppressed Patient Supports SARS-CoV-2 Viral Reservoir Hypothesis. Viruses 16 (2024). 10.3390/v16030342

9. Schmidt, H. et al. Adaptive evolution of SARS-CoV-2 during a persistent infection for 521 days in an immunocompromised patient. npj Genomic Medicine 10 (2025). 10.1038/s41525-025-00463-x

10. Li, Y. et al. SARS-CoV-2 viral clearance and evolution varies by type and severity of immunodeficiency. Science Translational Medicine 16 (2024). 10.1126/scitranslmed.adk1599

11. Marques, A. D. et al. SARS-CoV-2 evolution during prolonged infection in immunocompromised patients. mBio 15 (2024). 10.1128/mbio.00110-24

12. Clark, S. A. et al. SARS-CoV-2 evolution in an immunocompromised host reveals shared neutralization escape mechanisms. Cell 184, 2605–2617.e2618 (2021). 10.1016/j.cell.2021.03.027

13. Ko, K. K. K. et al. Emergence of SARS-CoV-2 Spike Mutations during Prolonged Infection in Immunocompromised Hosts. Microbiology Spectrum 10 (2022). 10.1128/spectrum.00791-22

14. Corey, L. et al. SARS-CoV-2 Variants in Patients with Immunosuppression. New England Journal of Medicine 385, 562–566 (2021). 10.1056/NEJMsb2104756

15. Carabelli, A. M. et al. SARS-CoV-2 variant biology: immune escape, transmission and fitness. Nature Reviews Microbiology (2023). 10.1038/s41579-022-00841-7

16. Kumata, R. & Sasaki, A. Antigenic escape is accelerated by the presence of immunocompromised hosts. Proceedings of the Royal Society B: Biological Sciences 289 (2022). 10.1098/rspb.2022.1437

17. Van Egeren, D. et al. Controlling long-term SARS-CoV-2 infections can slow viral evolution and reduce the risk of treatment failure. Scientific Reports 11 (2021). 10.1038/s41598-021-02148-8

18. Ghafari, M., Liu, Q., Dhillon, A., Katzourakis, A.& Weissman, D.B. Investigating the evolutionary origins of the first three SARS-CoV-2 variants of concern. Frontiers in Virology 2 (2022). 10.3389/fviro.2022.942555

19. Smith, C. A. & Ashby, B. Antigenic evolution of SARS-CoV-2 in immunocompromised hosts. Evolution, Medicine, and Public Health 11, 90–100 (2023). 10.1093/emph/eoac037

20. Coats, A., Wang, Y.R.& Koelle, K. Immune pressure is key to understanding observed patterns of respiratory virus evolution in prolonged infections. Virus Evolution 11 (2025). 10.1093/ve/veaf054

21. Louten, J. in Essential Human Virology p71–92 (2016).

22. Jenner, A. L. et al. COVID-19 virtual patient cohort suggests immune mechanisms driving disease outcomes. PLOS Pathogens 17, e1009753–e1009753 (2021). 10.1371/journal.ppat.1009753

23. Smith, A. P., Moquin, D. J., Bernhauerova, V. & Smith, A. M. Influenza virus infection model with density dependence supports biphasic viral decay. Frontiers in Microbiology 9, 1554–1554 (2018).

24. Deng, X., Farhang-Sardroodi, S.& Craig, M. Predicting age-related determinants of heterogeneous outcomes to COVID-19 mRNA vaccines through mathematical modelling. medRxiv, 1–38 (2025). 10.1101/2025.02.14.25322308

25. Owens, K., Esmaeili, S. C Schiffer, J.T. Heterogeneous SARS-CoV-2 kinetics due to variable timing and intensity of immune responses. JCI Insight 9 (2024). 10.1172/jci.insight.176286

26. Ciupe, S. M., Ribeiro, R. M., Nelson, P. W., Dusheiko, G. & Perelson, A. S. The role of cells refractory to productive infection in acute hepatitis B viral dynamics. Proceedings of the National Academy of Sciences 104, 5050–5055 (2007). 10.1073/pnas.0603626104

27. Iyaniwura, S. A. et al. The kinetics of SARS-CoV-2 infection based on a human challenge study. Proceedings of the National Academy of Sciences 121 (2024). 10.1073/pnas.2406303121

28. Samuel, C.E.& Knutson, G.S. Mechanism of interferon action. Kinetics of decay of the antiviral state and protein phosphorylation in mouse fibroblasts treated with natural and cloned interferons. Journal of Biological Chemistry 257, 11796–11801 (1982). 10.1016/s0021-9258(18)33834-1

29. Hayden, R. T., Wolk, D. M., Carroll, K. C. & Tang, Y.-W. Ch. 1, (2016).

30. Huang, B. et al. Interferon response and profiling of interferon response genes in peripheral blood of vaccine-naive COVID-19 patients. Frontiers in Immunology 14 (2024). 10.3389/fimmu.2023.1315602

31. Liu, B. C., Sarhan, J.& Poltorak, A. Host-Intrinsic Interferon Status in Infection and Immunity. Trends in Molecular Medicine 24, 658–668 (2018). 10.1016/j.molmed.2018.06.004

32. Fuchs, S. Y. Hope and Fear for Interferon: The Receptor-Centric Outlook on the Future of Interferon Therapy. Journal of Interferon & Cytokine Research 33, 211–225 (2013). 10.1089/jir.2012.0117

33. Sigal, A., Neher, R.A.& Lessells, R.J. The consequences of SARS-CoV-2 within-host persistence. Nature Reviews Microbiology 23, 288–302 (2024). 10.1038/s41579-024-01125-y

34. Smith, T.& Cunningham-Rundles, C. Primary B-cell immunodeficiencies. Human Immunology 80, 351–362 (2019). 10.1016/j.humimm.2018.10.015

35. Beachboard, D.C.& Horner, S.M. Innate immune evasion strategies of DNA and RNA viruses. Current Opinion in Microbiology 32, 113–119 (2016). 10.1016/j.mib.2016.05.015

36. Minkoff, J.M.& tenOever, B. Innate immune evasion strategies of SARS-CoV-2. Nature Reviews Microbiology (2023). 10.1038/s41579-022-00839-1

37. Randall, R. E. & Goodbourn, S. Interferons and viruses: an interplay between induction, signalling, antiviral responses and virus countermeasures. Journal of General Virology 89, 1–47 (2008). 10.1099/vir.0.83391-0

38. Samuel, C. E. Antiviral Actions of Interferons. Clinical Microbiology Reviews 14, 778– 809 (2001). 10.1128/cmr.14.4.778-809.2001

39. Day, T., Kennedy, D. A., Read, A. F. & Gandon, S. Pathogen evolution during vaccination campaigns. PLOS Biology 20, e3001804 (2022). 10.1371/journal.pbio.3001804

40. Lion, S. Class Structure, Demography, and Selection: Reproductive-Value Weighting in Nonequilibrium, Polymorphic Populations. The American Naturalist 191, 620–637 (2018). 10.1086/696976

41. Otto, S. P. & Day, T. A biologist’s guide to mathematical modeling in ecology and evolution. (Princeton University Press, 2007).

42. Taylor, P. D. Allele-Frequency Change in a Class-Structured Population. Am Nat 135, 95–106 (1990). 10.1086/285034

43. Laing, A. G. et al. A dynamic COVID-19 immune signature includes associations with poor prognosis. Nature Medicine 26, 1623–1635 (2020). 10.1038/s41591-020-1038-6

44. Liu, T. et al. The role of interleukin-6 in monitoring severe case of coronavirus disease 2019. EMBO Mol Med 12, e12421 (2020). 10.15252/emmm.202012421

45. Slatkin, M. Linkage disequilibrium — understanding the evolutionary past and mapping the medical future. Nature Reviews Genetics 9, 477–485 (2008). 10.1038/nrg2361

46. Felsenstein, J. The Effect of Linkage on Directional Selection. Genetics 52, 349–363 (1965). 10.1093/genetics/52.2.349

47. de Visser, J. A. G. M., Cooper, T. F. & Elena, S. F. The causes of epistasis. Proceedings of the Royal Society B: Biological Sciences 278, 3617–3624 (2011). 10.1098/rspb.2011.1537

48. McLeod, D. V. & Gandon, S. Understanding the evolution of multiple drug resistance in structured populations. eLife 10 (2021). 10.7554/eLife.65645

49. Holmes, E. C. The Emergence and Evolution of SARS-CoV-2. Annual Review of Virology 11, 21–42 (2024). 10.1146/annurev-virology-093022-013037

50. Cele, S. et al. SARS-CoV-2 prolonged infection during advanced HIV disease evolves extensive immune escape. Cell Host & Microbe 30, 154–162.e155 (2022). 10.1016/j.chom.2022.01.005

51. Chen, L. et al. Emergence of Multiple SARS-CoV-2 Antibody Escape Variants in an Immunocompromised Host Undergoing Convalescent Plasma Treatment. mSphere 6 (2021). 10.1128/mSphere.00480-21

52. Thorne, L. G. et al. Evolution of enhanced innate immune evasion by SARS-CoV-2. Nature 602, 487–495 (2021). 10.1038/s41586-021-04352-y

53. Harari, S. et al. Drivers of adaptive evolution during chronic SARS-CoV-2 infections. Nature Medicine 28, 1501–1508 (2022). 10.1038/s41591-022-01882-4

54. Eden, J.-S. et al. Persistent infections in immunocompromised hosts are rarely sources of new pathogen variants. Virus Evolution 3 (2017). 10.1093/ve/vex018

55. Watanabe, S., Alexander, M., Misharin, A. V. & Budinger, G. R. S. The role of macrophages in the resolution of inflammation. Journal of Clinical Investigation 129, 2619–2628 (2019). 10.1172/jci124615

56. Arish, M.& Sun, J. Monocyte and macrophage function in respiratory viral infections. Animal Diseases 3 (2023). 10.1186/s44149-023-00095-7

57. Takahashi, T. et al. Sex differences in immune responses that underlie COVID-19 disease outcomes. Nature 588, 315–320 (2020). 10.1038/s41586-020-2700-3

58. Sokoya, T., Steel, H. C., Nieuwoudt, M.& Rossouw, T.M. HIV as a Cause of Immune Activation and Immunosenescence. Mediators of Inflammation 2017, 1–16 (2017). 10.1155/2017/6825493

59. Touizer, E. et al. Failure to seroconvert after two doses of BNT162b2 SARS-CoV-2 vaccine in a patient with uncontrolled HIV. The Lancet HIV 8, e317–e318 (2021). 10.1016/s2352-3018(21)00099-0

60. Mancuso, S., Mattana, M., Carlisi, M., Santoro, M.& Siragusa, S. Effects of B-Cell Lymphoma on the Immune System and Immune Recovery after Treatment: The Paradigm of Targeted Therapy. International Journal of Molecular Sciences 23 (2022). 10.3390/ijms23063368

61. Nassef Kadry Naguib Roufaiel, M., Wells, J.W.& Steptoe, R.J. Impaired T-Cell Function in B-Cell Lymphoma: A Direct Consequence of Events at the Immunological Synapse? Frontiers in Immunology 6 (2015). 10.3389/fimmu.2015.00258

62. Luria, S.E.& Delbrück, M. Mutations of Bacteria from Virus Sensitivity to Virus Resistance. Genetics 28, 491–511 (1943). 10.1093/genetics/28.6.491

63. Otto, S. P. et al. The origins and potential future of SARS-CoV-2 variants of concern in the evolving COVID-19 pandemic. Current Biology 31, R918–R929 (2021). 10.1016/j.cub.2021.06.049

64. Myers, M. A. et al. Dynamically linking influenza virus infection kinetics, lung injury, inflammation, and disease severity. eLife 10 (2021). 10.7554/eLife.68864

65. Smith, A. M. & Perelson, A. S. Influenza A virus infection kinetics: quantitative data and models. Wiley Interdisciplinary Reviews. Systems Biology and Medicine 3, 429– 445 (2011). 10.1002/wsbm.129

66. Rochman, I., Paul, W. E. & Ben-Sasson, S. Z. IL-6 Increases Primed Cell Expansion and Survival. The Journal of Immunology 174, 4761–4767 (2005). 10.4049/jimmunol.174.8.4761

67. Soumelis, V. et al. Depletion of circulating natural type 1 interferon-producing cells in HIV-infected AIDS patients. Blood 98, 906–912 (2001). 10.1182/blood.V98.4.906

68. Hart, L. et al. Burden of chemotherapy-induced myelosuppression among patients with extensive-stage small cell lung cancer: A retrospective study from community oncology practices. Cancer Medicine 12, 10020–10030 (2023). 10.1002/cam4.5738

69. Cassidy, T., Humphries, A. R., Craig, M.& Mackey, M.C. Characterizing Chemotherapy-Induced Neutropenia and Monocytopenia Through Mathematical Modelling. Bulletin of Mathematical Biology 82, 104–104 (2020). 10.1007/s11538-020-00777-0

70. Holt, C. D. Overview of Immunosuppressive Therapy in Solid Organ Transplantation. Anesthesiology Clinics 35, 365–380 (2017). 10.1016/j.anclin.2017.04.001

71. Farhang-Sardroodi, S., Deng, X., Portet, S., Arino, J.& Craig, M. Insights into B cell and antibody kinetics against SARS-CoV-2 variants using mathematical modelling. bioRxiv (2023). 10.1101/2023.11.10.566587

72. Ma, Y., Zhang, Y. & Zhu, L. Role of neutrophils in acute viral infection. Immunity, Inffammation and Disease G, 1186–1196 (2021). 10.1002/iid3.500

